# Linker-rigidified VHL homodimerizers convert degraders into stabilizers of non-ubiquitinable ternary complexes

**DOI:** 10.1101/2025.11.18.689129

**Authors:** Charlotte Crowe, Alessandra Salerno, Gajanan Sathe, Graham Marsh, Hannah Maple, Alessio Ciulli

## Abstract

Protein degraders recruit E3 ligases to targets for ubiquitination and subsequent proteasomal degradation. Efficient degradation typically correlates with long-lived ternary complexes that position target lysines for productive ubiquitination. Here, we determine a cryo-EM structure of the Cullin 2 RING VHL (CRL2^VHL^) ligase dimerized by the VHL homo-PROTAC CM11, wherein one CRL^VHL^ acts as the ligase and a second VHL as the neo-substrate. Guided by this structure, we design side-by-side, linker-rigidified VHL homodimerizers that bias the relative VHL orientation away from the E2∼Ub active site, contrasting the flexible, head-to-head PEG linkage of CM11. Biophysical binding and in vitro ubiquitination assays show that these compounds form stable, long-lived and compact ternary complexes that are incompatible with VHL cross-ubiquitination. In cells, the compounds stabilize VHL and concomitantly inhibit it to elevate HIF-1α levels. Thus, a stable ternary complex can be non-productive for ubiquitination, and linker architecture can reprogram degraders into stabilizers by controlling target ubiquitinability.

## Introduction

Targeted protein degradation (TPD) removes disease-relevant proteins by co-opting a ubiquitin E3 ligase to trigger ubiquitination and subsequent proteasomal degradation of non-native substrates. Bifunctional proteolysis-targeting chimeras (PROTACs) recruit an E3 ligase and a target protein via two respective ligands joined by a linker,^1,2^ whereas molecular glue degraders are typically monovalent and promote interactions between the protein partners.^3,4^ Recent work has revealed mechanisms spanning this continuum.^5,6^ Regardless of recruitment mode, protein degraders depend on E3-mediated ubiquitination of the captured neo-substrate.

Most TPD campaigns recruit Cullin RING E3 ligases (CRLs),^7^ with Cullin2 RING E3 ligase von Hippel-Lindau (CRL2^VHL^)^8–10^ and Cullin 4 RING E3 ligase cereblon (CRL4^CRBN^)^11–13^ being the most widely used. CRL2^VHL^ assembles from VHL, adaptor proteins EloB and EloC, the scaffold protein Cul2 and the RING protein Rbx1. Two major VHL isoforms exist: a ‘long’ pVHL30 (VHL_1-213_) isoform, and a ‘short’ pVHL19 (VHL_54-213_) isoform; the latter lacking the N-terminal acidic repeat domain,^14,15^ yet retaining core E3 ligase functions and localizing to both cytosol and nucleus.^16^ CRL2^VHL^ primary function is to target hypoxia-inducible factor alpha (HIF-α) subunits for oxygen-dependent proteolysis.^17^ Following significant advances in designing high-affinity ligands for the VHL substrate receptor,^18–22^ VHL has been broadly used and applied in PROTAC development campaigns for TPD.^10^

Until recently, most structural analyses of degrader ternary complexes focused on substrate receptors and adaptors,^5,9,12^ limiting insights into how full CRL assemblies position substrates for ubiquitination. Biochemical and cellular studies of degrader-driven ubiquitination have likewise been limited in resolution. Although ternary complex stability and half-life often correlate with degradation efficacy and selectivity,^9,23–29^ ubiquitination may still fail due to non-productive complex geometries or UPS resistance.^30,31^ Thus, defining how ternary complex orientation within the full E3 ligase dictates “ubiquitinability” is paramount. Recent full-complex studies have shown that the well-characterized PROTAC MZ1 (ref. ^32^) forms a highly cooperative and stable ternary complex with CRL2^VHL^ and the *neo*-substrate Brd4^BD^^2^, that drives rapid and profound degradation of Brd4,^9,25^ by positioning lysines of Brd4^BD2^ optimally for ubiquitination.^33,34^ However, the behavior of other degrader-induced ternary complexes within the complete CRL2^VHL^ remains incompletely characterized.

We were curious to explore the ternary complex mediated by CM11, a homoPROTAC which dimerizes and degrades VHL.^35^ The molecule, developed and disclosed in 2017 by Maniaci *et al.*, consists of two copies of VHL ligand VH032 (ref. ^20^) connected by a PEG_5_ linker. Our previous strategy aimed to form a 2:1 VHL:homoPROTAC assembly where CRL2^VHL^ acts as both the E2-Ub recruiting E3 ligase as well as the targeted substrate. We previously leveraged co-crystal structures of VHL bound to VH032 (ref. ^20^) and VH298 (ref. ^21^) (PDB codes 4W9H and 5LLI, respectively) to define left-hand (LHS, acetyl) and right-hand (RHS, phenyl) exit vectors for linker attachment. We synthesized (1) symmetric head-to-head VH032 homodimers with varying PEG lengths (PEG_3_-PEG_5_) from the LHS; (2) symmetric side-by-side homodimers from the RHS; (3) asymmetric head-to-side constructs.^35^ Only the LHS head-to-head series induced robust VHL self-degradation, with the PEG_5_-linked CM11 being the most effective, degrading pVHL30 in a time- and concentration-dependent manner and partially reducing pVHL19 and Cul2, thus resulting in only minimal increase in hydroxylated HIF-1α protein level. Biophysical assays showed a stable and cooperative 2:1 VHL:CM11 ternary complex (α ∼ 20) for both isoforms.^35^ By contrast, side-by-side and asymmetric designs were inactive, underscoring the importance of linkage geometry. These features, together with emerging principles of substrate “ubiquitinability” within full CRL2^VHL^ assemblies,^33^ motivated us to select CM11 as a model to interrogate further the relationships between ternary complex geometry and ubiquitinability.

## RESULTS

### Cryo-EM structure of CM11 bound to a dimer of CRL2^VHL^

From a structural point of view, the VHL protein (short isoform pVHL19, 18 kDa or long isoform pVHL30, 24 kDa) is the substrate receptor of the pentameric 150 kDa CRL2^VHL^ E3 ligase. The largest component of this assembly is Cullin 2 (Cul2, 87 kDa) which serves as a flexible scaffolding protein. VHL is recruited to Cul2 via the adaptor proteins Elongin C (EloC, 12 kDa) and Elongin B (EloB, 15 kDa).^8,36^ The RING box protein 1 (Rbx1, 12 kDa) enables the recruitment of E2 conjugating enzymes. These E2 proteins can either be ubiquitin-conjugating, which enables ubiquitination of substrates recruited by VHL, or NEDD8-conjugating, which triggers auto-neddylation of CRL2^VHL^ at the K689 residue of the Cul2 WHB domain.^37,38^ This latter modification has been shown to induce conformational rearrangements of Cullin-RING complexes, thereby facilitating the engagement of E2 ubiquitin-conjugating enzymes and thus activating the E3 ligase complex for ubiquitination.^39–41^

Our multiple attempts at co-crystallizing either CM11 or the analogous SH2, both symmetric head-to-head VHL-degrading homoPROTACs,^35,42^ when simultaneously bound in complex with two molecules of VHL-EloB-EloC (VCB), have to date remained unsuccessful, preventing structural determination by X-ray crystallography. We therefore turned to cryogenic electron microscopy (cryo-EM) to illuminate structural information on a 1:2 CM11:VHL assembly. Towards this goal, we expressed and purified the Cul2-Rbx1 scaffold component, and incubated it with VCB to form the pentameric complex VHL-EloC-EloB-Cul2-Rbx1, using the short pVHL19 isoform given that the N-terminal residues 1-54 are predicted to constitute an unstructured flexible tail. The fully-formed CRL2^VHL^ was *in vitro* neddylated with NEDD8 using APPBP1-UBA3 and UBE2M, and purified by size-exclusion chromatography to isolate pure NEDD8-CRL2^VHL^ (**Figure S1 A-C**).

We next set to perform cryo-EM analysis of the E3 ligase NEDD8-CRL2^VHL^ in complex with CM11. Samples of 0.05 mg/mL recombinant NEDD8-CRL2^VHL^ protein were mixed with CM11 in DMSO, incubated over ice for 10 minutes, then desalted through a spin desalting column. Presence of the ternary complex was confirmed by mass photometry (**Figure S1D**). The resulting sample was applied to copper Quantifoil R1.2/1.3 grids, which were plunge-frozen on a Vitrobot Mark IV and imaged on a Glacios 200 kV with Falcon 4i. Subsequent data collection of 7,216 movies and image analysis resulted in a 7 Å resolution structure (**Figure S2**). Despite the modest resolution, the 2D classes showed good distribution of orientations and a wide range of views (**Figure S2**) and the volumes were of sufficient quality and clarity to fit models of CRL2^VHL^ (**Figure 1A**).

**Figure 1.**
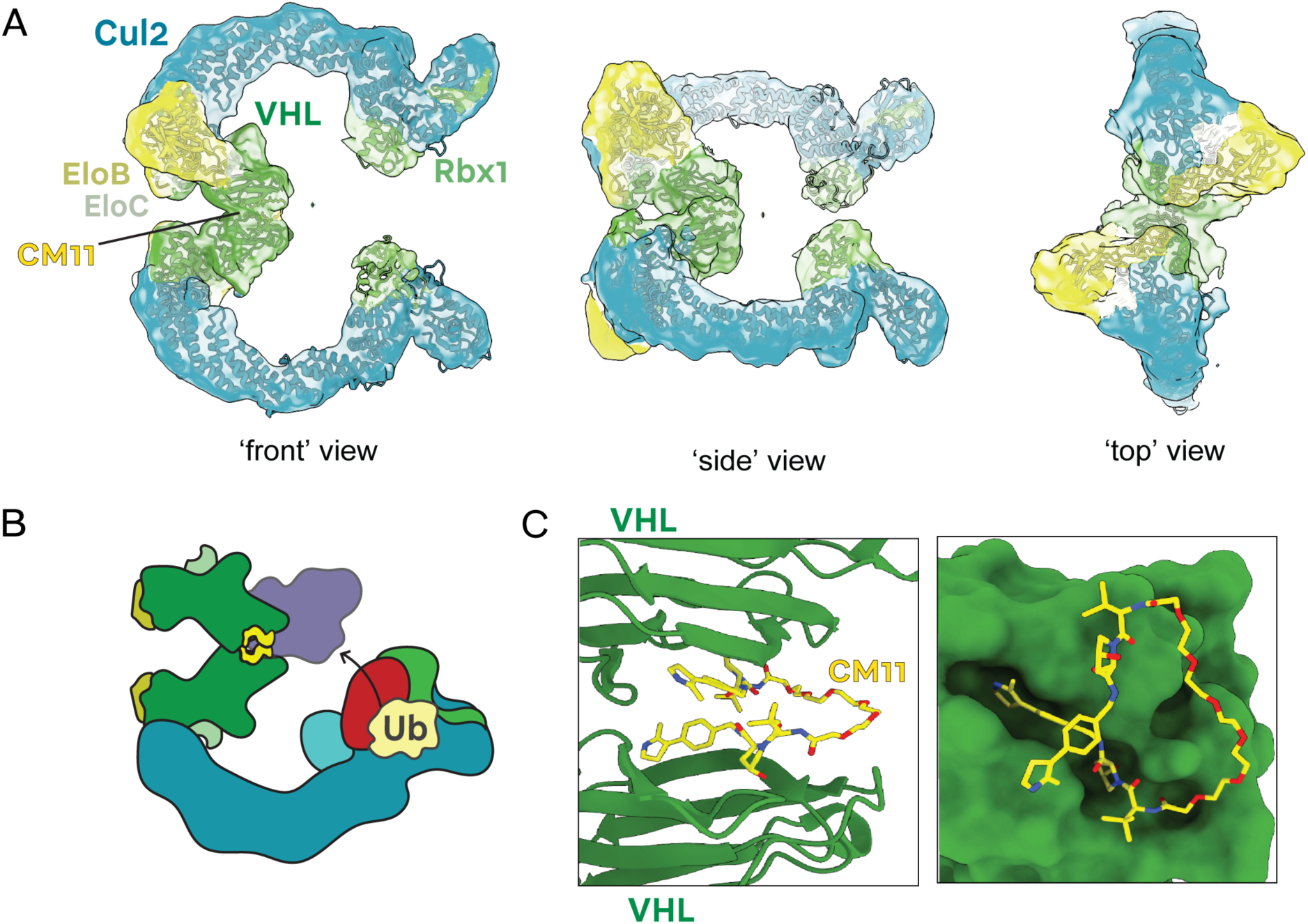
Cryo-EM analysis of CM11 in complex with two molecules of (NEDD8)-CRL2^VHL^. **(A)** Cryo-EM volume (opaque) with fitted model of (NEDD8)-CRL2^VHL^ dimerized by CM11 (ribbon diagram). The domains are colored: VHL (forest green), EloC (light green), EloB (khaki), Cul2 (turquoise), Rbx1 (lime green). Front, side and top views are shown. **(B)** Schematic illustrating the superimposition of the BRD4^BD2^-MZ1-(NEDD8)-CRL2^VHL^-UBE2R1-Ub structure^33^ with the cryo-EM model in this study, showing how BRD4^BD2^ (purple) is oriented by MZ1 towards the UBE2R1-Ub catalytic site, whereas VHL (forest green) is oriented by CM11 in the opposite direction. The domains are colored: BRD4^BD2^ (purple), NEDD8 (cyan), UBE2R1 (red) and ubiquitin (yellow). **(C)** Zoom in to the modelled CM11 homoPROTAC. Left: a front view is shown with the VHL-VHL interface of the fitted model (forest green, ribbon diagram). Right: a side view is shown of CM11 with a VHL monomer of fitted model (forest green, surface).

Our cryo-EM volumes provide a first opportunity to appreciate the overall geometry of NEDD8-CRL2^VHL^ dimerization induced by CM11. **Figure 1A** shows a ‘glueing’ of the VHL-VHL interface, consistently with the best resolved section of the cryo-EM 2D classes and 3D volumes in our analysis (**Figure S2**). This interface directs the overall cullin RING E3 into a ‘heart-shaped’ configuration, with VHL directed away from the cullin C-terminus and RING domain. This was a striking contrast to the previously published BRD4^BD2^-MZ1-(NEDD8)-CRL2^VHL^-UBE2R1-Ub cryo-EM structure,^33^ where MZ1 orients BRD4^BD2^ towards the UBE2R1-Ub catalytic site for ubiquitination (**Figure 1B**, **Figure S3**). We also noted that only three lysine residues are present on VHL: of these, one is inaccessible at the interface with Elongin C and Cullin 2, and the other two are located on alpha helices furthest from the dimerized Cullin C-termini, mapping to a non-ubiquitinable ‘dark face’^33^ on VHL as modelled in our structure. These observations suggested CM11 might mediate a less ubiquitinable ternary complex compared to MZ1. Given that CM11 indeed degrades VHL,^35^ we sought to rationalize how ubiquitin transfer could occur in this complex. Interestingly, at the 2D class level from the cryo-EM dataset, we observed some particle stacks which deviated from the 3D consensus volume, suggesting conformational heterogeneity in the sample and thus more opportunity for VHL ubiquitination than suggested by the structural model (**Figure S4**).

Modelling of CM11 at the VHL-VHL interface of the cryo-EM structure was possible by leveraging the numerous co-crystal structures of VHL ligands in complex with VCB, including the most relevant with bound VH032 (ref. ^20^). Our model shows how CM11 folds on itself to mediate VHL dimerization, and how the PEG_5_ linker must cover a long distance to allow for the two VH032 units to achieve their ‘crossed’ relative conformation (**Figure 1C**). These new insights into the VHL-VHL assembly enabled us to retrospectively rationalize the structure-activity relationship observed by Maniaci *et al.*,^35^ whereby degradation activity decreased with linker-shortening such that, in order of potency, CM11 (PEG_5_) >> CM10 (PEG_4_) > CM09 (PEG_3_) (**Figure S5A**). Indeed, both CM09 and CM10 exhibited minimal ternary complex formation, with the shorter linker in CM09 correlating with significantly reduced degradation of pVHL30, no degradation of pVHL19 or Cul2, and no increase in HIF-1α levels.^35^ The relative configuration of the VH032 ligands within the VHL binding pockets appeared to be enabled by the flexibility of the PEG_5_ linker, explaining why the shorter PEG linkers tested by Maniaci *et al*. were less effective, likely due to their limited ability to adopt the same stable ternary complex as CM11.

### Structure-based design of linker-optimized CRL2^VHL^ dimerizers

Given the structural information on the CM11-mediated CRL2^VHL^ ternary complex, we posited that we could use this structure to inspire and guide the rational design of linker-optimized dimerizers, by replacing the CM11 PEG_5_ linker region with more structurally favorable VHL ligand exit vectors and atom-efficient linkers. Owing to its increased binary affinity towards VHL and increased cell permeability, we selected the VH101 ligand (**Figure S5B**)^22^ instead of VH032 as a VHL ligand scaffold, and used the available co-crystal structures of VH101 in complex with the VCB monomer for modelling into the cryo-EM structure.^22^ We examined 15 sharpened and unsharpened cryo-EM maps from independent refinements with both C1 and C2 symmetry and performed rigid-body docking of the VHL subunit of the co-crystal structures into each of these maps to obtain a distribution of relative VHL-VH101 poses (**Figure S6A**). By aligning all the poses on a single VHL-VH101 monomer on one side of the model (**Figure S6B**), we obtained a narrow range of VH101-VH101 relative poses (**Figure S6C**). From visual inspection and manual modelling, the phenyl-benzylic connection between two distinct VH101 units was identified as a structurally favorable and synthetically tractable connection point, in an unprecedented ‘side-to-side’ configuration as opposed to the traditional ‘head-to-head’ connection often employed by linear PROTACs. The required phenyl-benzylic distance could be bridged by a short 2-atom linker with sensible bond angles (**Figure S6D**). We thus leveraged this structural information to design and synthesize three new ‘side-to-side’ dimeric VHL compounds (**Figure 2B**).

**Figure 2.**
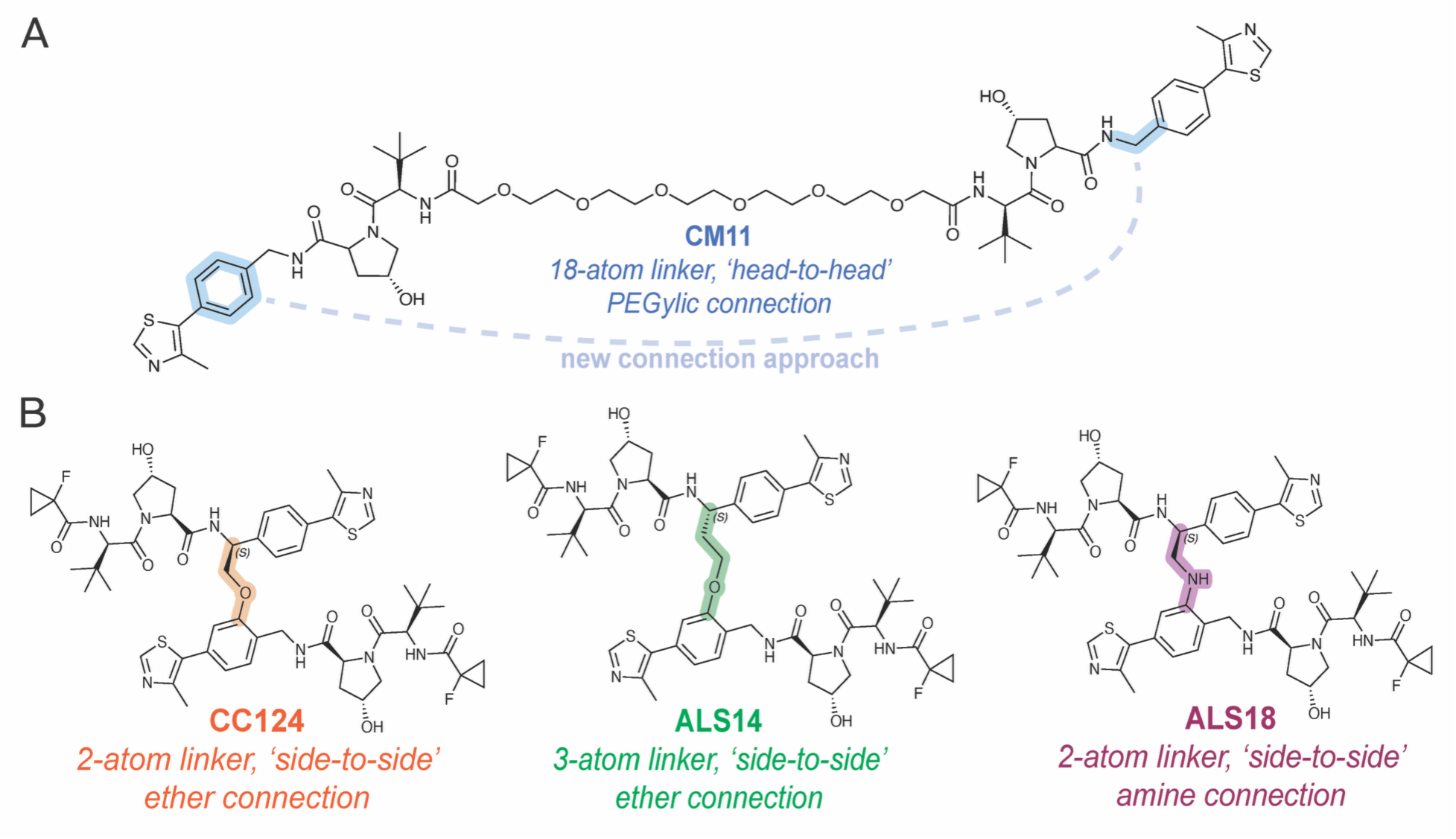
Design of new compounds in this study. **(A)** Chemical structure of the homoPROTAC CM11 with its flexible PEG5 linker. The proposed connection approach for this study is shown in blue. **(B)** New side-by-side compounds designed in this study, with the new phenyl-benzylic connection shown in orange, green and purple for CC124, ALS14 and ALS18 respectively.

Given the commercial availability of VH101-phenol, we reasoned that the phenolic moiety could serve as an exit vector for the phenyl connection. CC124 was thus obtained via a phenolic methyl ether bridge from the phenolic to the benzylic position of another VH101 scaffold. Using the same ‘side-to-side’ connection strategy, ALS14 was designed and synthesized to modulate linker length and enhance molecular flexibility by incorporating a three-atom phenolic ethyl ether linker. Finally, to exploit potential differences in intra-molecular hydrogen-bonding, we synthesized a third compound, ALS18, featuring a short phenylamino methyl amine using a custom ‘VH101 aniline’ building block. Achieving these compounds required redesign of synthetic routes, described in detail in the Supplementary Materials, **Schemes S1-3**.

### CRL2^VHL^ dimerizers engage VHL and form a long-lived ternary complex

The newly synthesized compounds were profiled for binary VHL engagement and ternary complex formation using the heliX^43^ technology. We conjugated 48-nucleotide long single-stranded DNA (ssDNA) to recombinant VCB protein and purified this by ion-exchange chromatography for use on the heliX^®^ biosensor. For binary binding, we immobilized ssDNA-VCB to the surface of a heliX^®^ biochip and used the heliX^®^ biosensor operating in fluorescence proximity sensing (FPS) mode^43^ to measure PROTAC binary binding with the heliX^®^ adapter strands (**Figure S7**). We also measured *in vitro* ternary complex formation by Förster Resonance Energy Transfer (FRET) using the heliX^®^ Y-structure,^43^ where both the arms of the Y-structure were functionalized with VCB (**Figure 3A** and **S8**).

**Figure 3.**
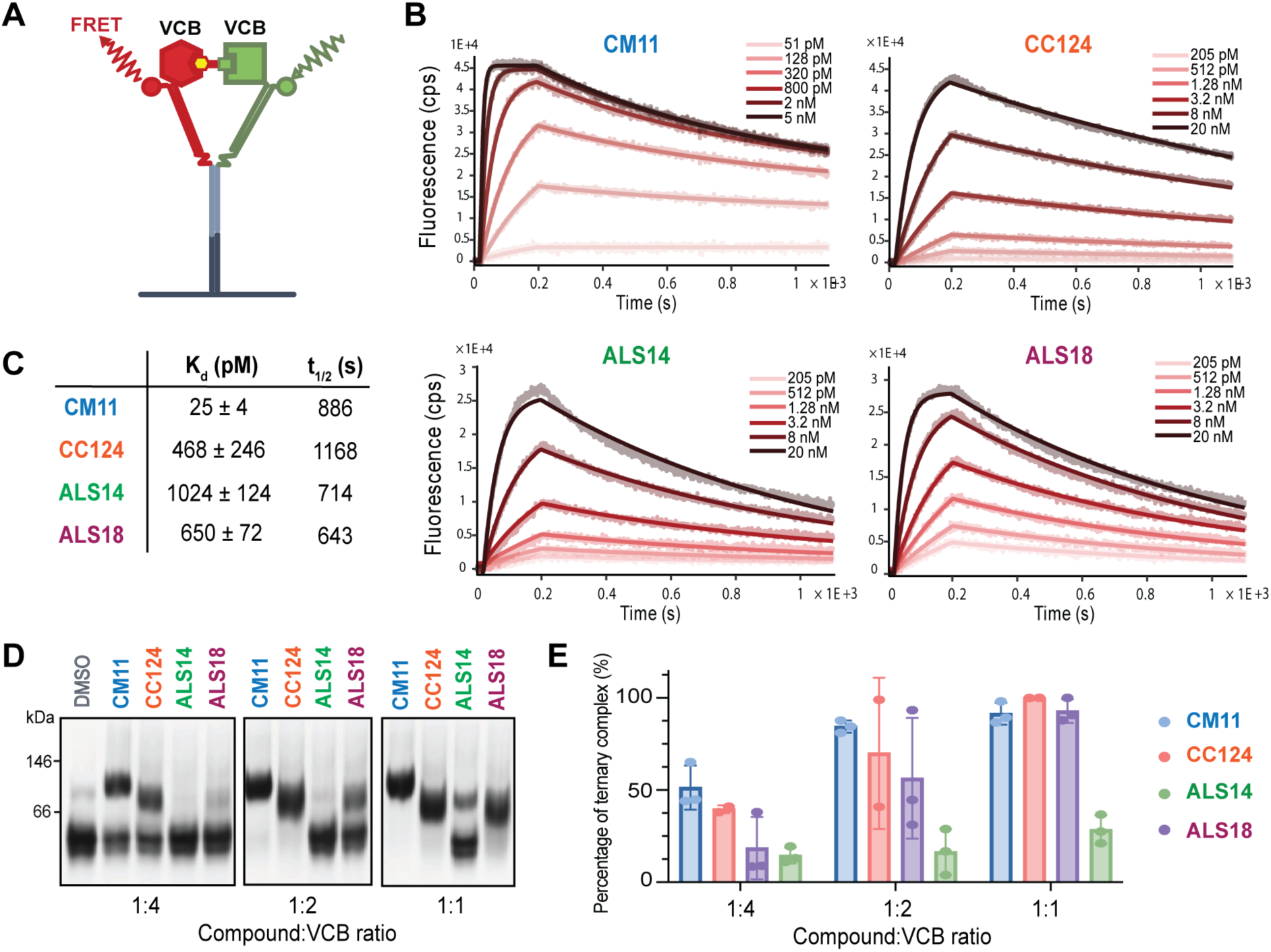
The homoPROTAC CM11 and side-by-side dimeric CC124, ALS14 and ALS18 compounds all form ternary complexes with VHL. **(A)** Schematic illustrating the assay set-up using the heliX Y-structure DNA strands and the emission of a FRET signal. **(B)** The heliX biosensor Y-structure measurements showed ternary complex formation for CM11 and the newly synthesized compounds. **(C)** Quantification of the ternary complex kinetic parameters, the error bars are shown as the standard deviation for *n* = 3 replicates. **(D)** EMSA showing the dimerisation of VCB by CM11, CC124, ALS14 and ALS18. **(E)** Quantification of the in-solution ternary complex formation resolved by EMSA, the error bars are shown as the standard deviation for *n* = 3 replicates.

At binary level, CM11 displayed the fastest rate of association with VCB at 158 × 10^5^ M^-1^ s^-1^, 4- to 5-fold faster than CC124, ALS14 and ALS18 (**Table S2**). This faster on-rate for 1:1 binding may be due to the linear CM11 compound being more flexible and less sterically constrained to engage with its binding partner. CM11 also had the fastest binary off-rate compared to our three new dimeric compounds, consistent with the lower binding affinity and faster off-rates of its VH032 ligand relative to VH101, as previously shown by us using SPR kinetics data with the monomeric VHL ligands.^22^

The measurement of ternary complex formation kinetics revealed a 30- to 18-fold faster association rate for CM11 as compared to CC124, ALS14 and ALS18, but similar dissociation rates, leading to a low *K*_d_ of 25 (±4) pM for VHL-CM11-VHL, and higher *K*_d_ values of 468 (±248) pM, 1024 (±124) pM and 650 (±72) pM for CC124, ALS14 and ALS18 complexes, respectively (**Figure 3B** and **C**). These binding affinities indicated that CM11, with its long linear PEG_5_ linker, is more apt at engaging both molecules of VHL at either end of the heliX Y-structure arms, likely because it is capable of ‘stretching out’ to reach the VBC molecules before folding into its preferred conformation, creating a tight VHL:VHL interface as shown in our cryo-EM analysis. By contrast, the geometrically constrained CC124, ALS14 and ALS18 compounds lack the ability to extend towards a second VHL unit, and once bound to the first VHL substrate, likely rely on a random interaction event where the other VHL monomer enters into close proximity with the preformed protein:compound binary species to form a ternary complex. Despite this obstacle to ternary association, the resulting ternary complexes were long-lived (with incomplete dissociation observed after 15 minutes of dissociation time in the assay) and had comparable longevity to the CM11-mediated complex.

To further understand the behaviors of this system, we sought to perform an in-solution assay, orthogonal to the heliX biosensor method. We carried out an electrophoresis mobility shift assay (EMSA) using a native gel, previously established by Diehl *et al.* and Salerno *et al.* for VHL homoPROTAC systems.^42,44^ We pre-incubated recombinant VCB protein with the bivalent dimerizer compounds or monovalent VH298 to assess the extent of VHL dimerization in solution (**Figure 3D**). In the EMSA assay, ternary complex formation is generally indicated by an upward shift in the protein band on the gel, in agreement with the higher molecular weight of the complex. Quantification of the proportion of VCB monomer versus dimer at three ratios of compound:VCB (1:4, 1:2, 1:1) showed good dimerization for CM11, CC124 and ALS18, and a lower proportion of dimerized VBC by ALS14 (**Figure 3E**), consistent with the ranking of *K*_d_ values calculated from the heliX Y-structure experiments. Interestingly, we noticed that the migration level of the protein bands through the native gel was slightly different for the different ternary complexes (**Figure S8**). Due to the lower proportion of ternary complex formation with ALS14, we compared CM11/ALS14/ALS18 using the same ratio but with a 10-fold dilution of VCB in an attempt to push dimerisation to completion. Given that migration of protein species through a native gel is governed by complex charge, size and shape, these results indicated a difference in the overall shape or conformation of the complexes. We observed a slight difference in the up-shifted migrated bands between CM11 and CC124, and three different band migrations between CM11, ALS14 and ALS18. We hypothesized that any major differences in overall relative geometry and orientation of the ternary complexes would likely result from the design of the dimeric compounds, considering their different conjugation vectors and linker lengths. Specifically, all the side-by-side dimeric compounds aligned to a more compact structure, with the 2-atom linkers (CC124 and ALS18) showing the greatest migration, and the 3-atom linker (ALS14) displaying intermediate migration. In contrast, the head-to-head PEG_5_ linker (CM11) exhibited the least migration indicative of a more heterogeneous assembly sampling wider space. We thus postulate that the more flexible CM11 is capable of mediating a larger range of conformations, consistent with our observations of heterogeneity in cryo-EM 2D classes, as opposed to the highly constrained compounds CC124 and ALS18 where the VCB-VCB dimer is likely locked into a single, more compact pose.

Next, to verify for engagement in a cellular milieu, we performed a NanoBRET cellular VHL engagement assay. Our results confirmed that CC124 and ALS18 engaged VHL in digitonin-permeabilized cells to a similar extent to CM11, while ALS14 showed a right-shifted curve, indicating weaker VHL engagement (**Figure S10**). This is consistent with reduced VHL association by ALS14 in solution by EMSA and on the biosensor surface by heliX. With this in mind, we proceeded with further biochemical and cellular evaluation.

### Linker rigidification of homo-dimerizers abolishes VHL ubiquitination

Given their stable ternary complexes, all three structure-based designed homo-dimerizers CC124, ALS14 and ALS18 were subsequently evaluated for their ability to mediate ubiquitination. We performed *in vitro* ubiquitination assays using recombinantly expressed and purified non-neddylated CRL2^VHL^ and NEDD8-CRL2^VHL^. Both forms of CRL2^VHL^ possessed the short isoform pVHL19. Although both the short (pVHL19) and long (pVHL30) isoforms are present in cell, these differ by the N-terminal residues 1-54, which are predicted to constitute an unstructured flexible tail. Since residues 1-54 do not contain any lysine residues, we were confident that the pVHL19-containing CRL2^VHL^ would constitute a representative species for *in vitro* ubiquitination study. We elected to use UBE2D2 and UBE2R1 as representative E2s from the UBE2D and UBE2R families, owing to their ability to work with CRL2 and ubiquitinate substrates.^45–47^ We resolved the protein species by SDS-PAGE, then visualized VHL (**Figure 4A** and **S11**) and Cul2 (**Figure 4B** and **S12**) ubiquitination by western blot, and confirmed ubiquitination of these species using an Alexa Fluor 488-labelled ubiquitin in the reaction and fluorescently imaging at 488 nm (**Figure S13**).

**Figure 4.**
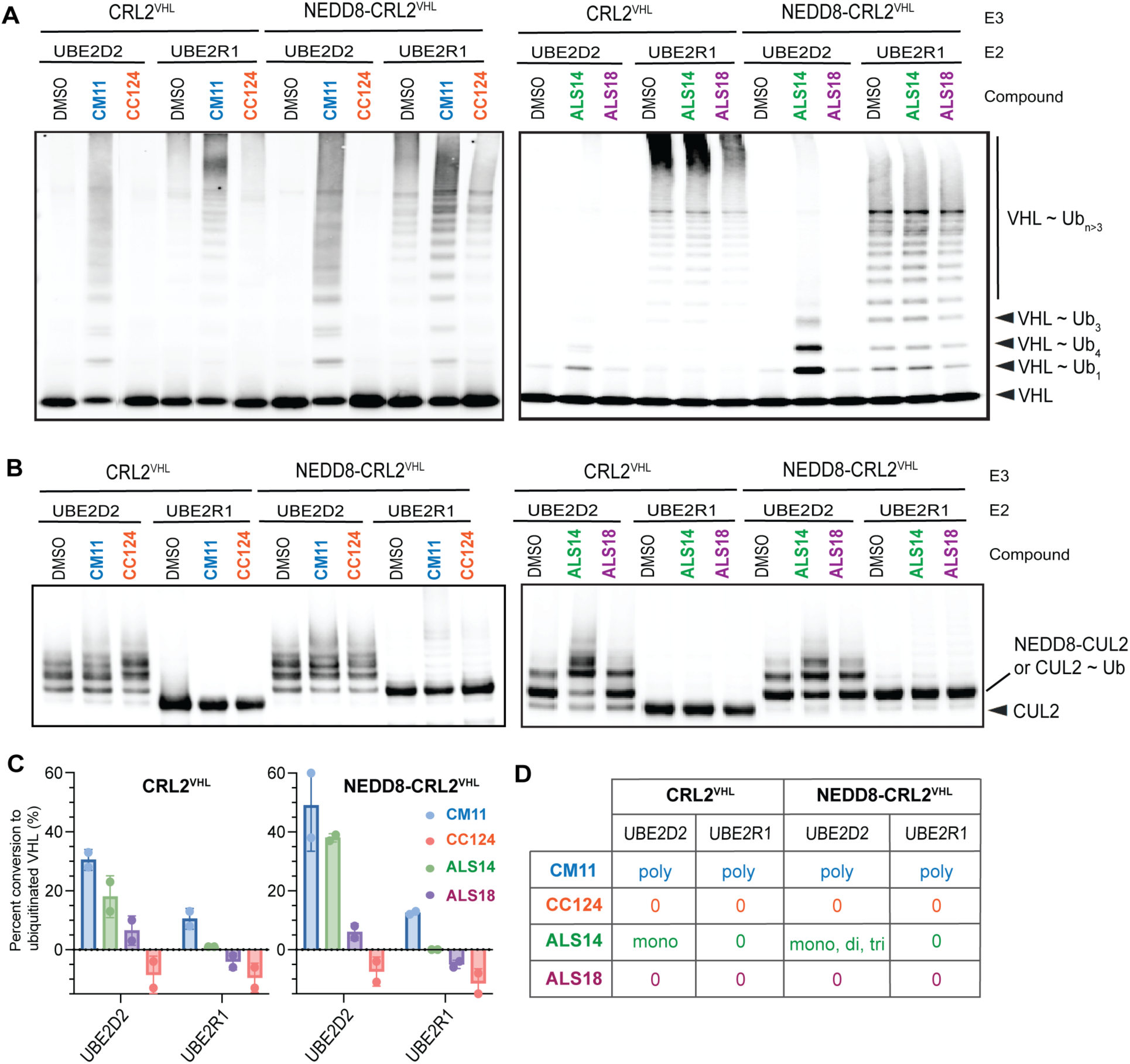
Side-by-side constrained VHL dimerizers CC124 and ALS18 do not induce VHL ubiquitination. **(A)** Representative Western Blot against VHL. (**B)** Representative Western Blot against Cul2. (**C)** Quantification of conversion from non-ubiquitinated VHL to ubiquitinated VHL for each dimerizer, relative to DMSO. The bars represent the mean of *n* = 2 biological replicates and the error bars correspond to the standard error of the mean. (**D)** Summary of VHL ubiquitination multiplicity for each E3 species, E2, and dimerizer as compared to DMSO. ‘Mono’, ‘di’, ‘tri’ and ‘poly’ refer to mono-, di-, tri- and poly-ubiquitinated VHL.

With UBE2D2, only CM11 and ALS14 mediated more ubiquitination on VHL than the DMSO-only control (**Figure 4A and C**). This effect was more pronounced with NEDD8-CRL2^VHL^ than with non-neddylated CRL2^VHL^, in line with the fact that neddylation has an activating effect on cullin RING E3s – upon neddylation, the contacts between the Cul2 WHB domain and Rbx1 are abrogated, ‘unlocking’ the cullin RING for E2 recruitment and catalysis.^39–41^ The increased linker flexibilities for CM11 (PEG_5_ linker) and ALS14 (3-atom linker) over CC124 and ALS18 (both 2-atom linkers), coupled with Cullin neddylation, enabled higher freedom of motion, and could rationalize why VHL can be ubiquitinated by CM11 and ALS14, but not by the highly constrained CC124 and ALS18 dimerizers.

Different ubiquitination patterns on VHL were observed when comparing CM11- and ALS14-mediated ubiquitination with UBE2D2 (**Figure 4D**). CM11 mediated monoubiquitination of VHL and also built polyubiquitin chains, as shown by the appearance of higher molecular weight species in the VHL western blot constituting a characteristic ‘polyubiquitin smear’ (**Figure 4A**). In contrast, the VHL western blot revealed that ALS14 treatment generated a pool of lower molecular weight monoubiquitinated VHL, with some less prominent di- and tri-ubiquitinated VHL bands visible. Given that CM11 should confer a higher degree of ternary complex flexibility, one could postulate that 1) multiple lysines on VHL are ubiquitinatable with CM11, and the species gets multi-monoubiquitinated, 2) polyubiquitin chains can be extended and the ternary complex is flexible enough to adapt to the steric bulk of many ubiquitination events, or a combination of 1) and 2). On the other hand, with ALS14, an accumulation of mono-ubiquitinated VHL was observed, suggesting the more constrained ternary complex is unable to access many lysines and/or cannot adapt to longer ubiquitin chains.

Using UBE2R1 in the reaction, only CM11 was able to ubiquitinate VHL more than the DMSO control (**Figure 4A and C**). UBE2R1 is known to be apt at extending polyubiquitin chains, but several studies have recently shown that UBE2R1 is also capable of installing the first ubiquitin on geometrically optimal substrates.^33,34^ Here, it appeared that VHL was not a geometrically optimal substrate for *cis*- or *trans*-ubiquitination, and only the high degree of flexibility conferred by the PEG_5_ linker, coupled with the inherent flexibility of the Cullin 2 scaffold, allows VHL to still be ubiquitinated in the presence of CM11.

With UBE2D2, monoubiquitination on non-neddylated Cullin 2 was observed with DMSO and all dimerizers in this study (**Figure 4B**). Ubiquitination independent of dimerizer presence is likely due to autoubiquitination at the WHB domain Lys689 which would otherwise be modified with NEDD8 (and we have indeed observed in other systems that Cullin 2 can be auto-ubiquitinated in the presence of UBE2D E2s and in the absence of NEDD8).^33^ By contrast, when NEDD8-CRL2^VHL^ was used in the assay in combination with UBE2D2, there was less conversion from non-ubiquitinated to ubiquitinated Cul2 (**Figure 4B**), likely due to the Lys689 site being already modified with NEDD8. Overall, quantification of Cullin 2 ubiquitination was more challenging due to dimerizer-independent auto-ubiquitination. We did however observe a slightly higher Cullin 2 ubiquitination in the presence of UBE2D2, namely for CM11, CC124 and ALS14 as compared with DMSO. Furthermore, for ALS14 some higher molecular weight species could be clearly observed by Cul2 immunoblot (**Figure 4B**) and at 488 nm (**Figure S13**), indicating that the ternary complex could be locked in such a way that allows for Cul2 to be better positioned for ubiquitination over the other dimerizer compounds. Using UBE2R1 alone, ubiquitination of Cul2 was either very low or undetectable for all dimerizers.

### Side-by-side homo-dimerizers stabilize and inhibit VHL in cells

Given the largely reduced or absence of VHL ubiquitination with our newly-designed dimerizers, we set to evaluate, by western blot analysis, their activity towards VHL and Cul2 protein levels after compound treatment for 24 hours (**Figure 5**) and 4 hours (**Figure S14**) in HEK293 cells. The degradation efficiency was compared to that of the reference homoPROTAC, CM11 and DMSO.

**Figure 5.**
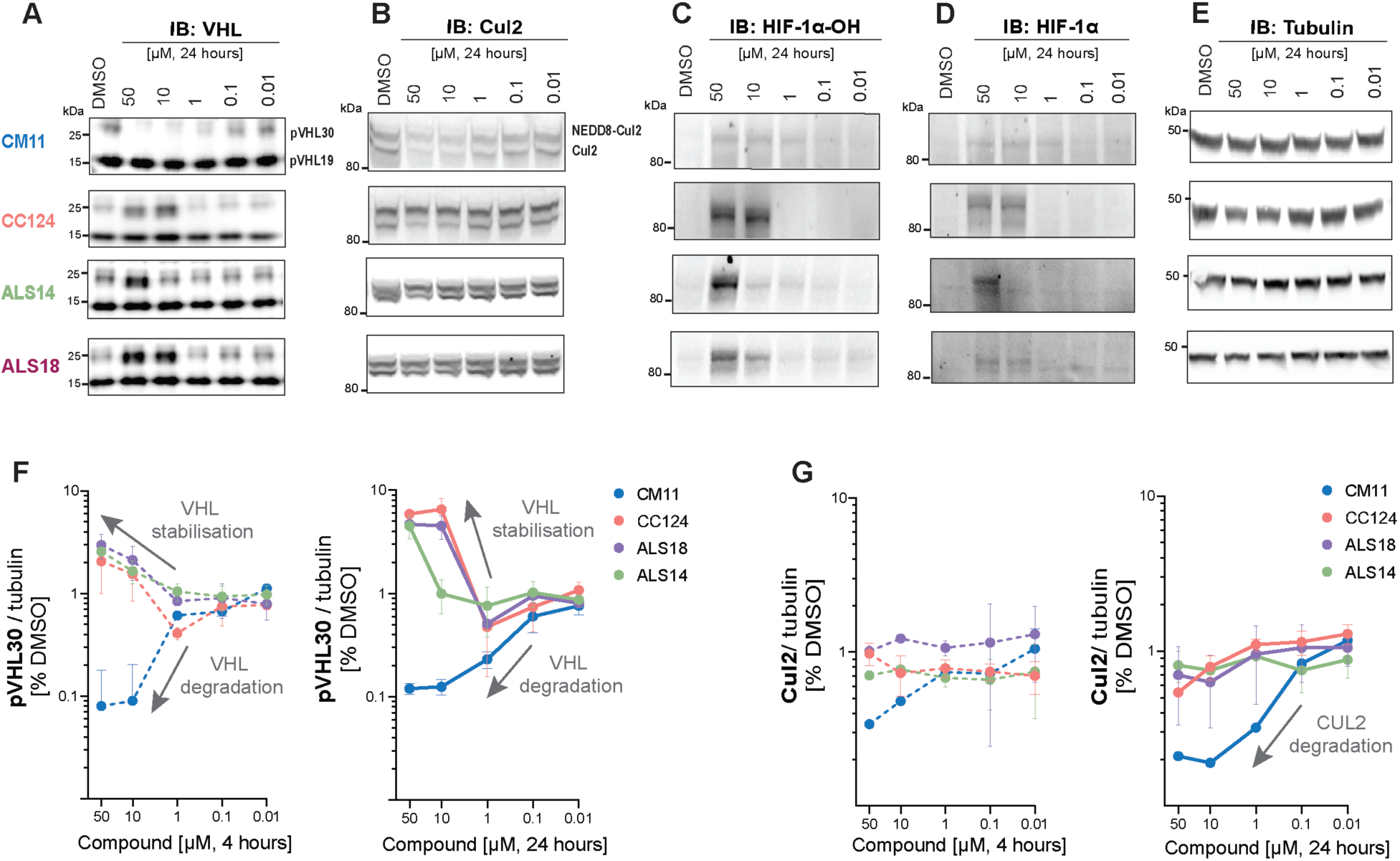
VHL dimerizers stabilize and inhibit VHL in cells. **(A-E)** Representative immunoblot of dose-dependent effects of compounds CM11, CC124, ALS14 and ALS18 on VHL, Cul2, HIF-1α-OH and HIF-1α, respectively in HEK293 cells (representative of *n* = 2 biological replicates, at indicated concentrations, 24 h) **(F)** Quantification of pVHL30 after 4 h and 24 h. **(G)** Quantification of Cul2 after 4 h and 24 h. The protein level/tubulin ratios were normalized to the average of the DMSO (100 %). Data represent the mean of *n* = 2 biological replicates and the error bars correspond to the standard error of the mean.

While CM11 demonstrated rapid and robust degradation of the pVHL30 isoform at both time points (**Figures 5A**, **5B** and **S14**), none of the newly synthesized compounds induced observable degradation of either of the VHL isoforms, consistent with the abrogation of ubiquitination we observed *in vitro*. Only a modest reduction in pVHL30 levels was observed with the 2-atom phenolic ether CC124 and the phenylamino ALS18 at 1 µM, mainly after 24 hours of treatment (**Figure 5B**). At concentrations higher than 1 µM, instead of mediating increased degradation, all three compounds CC124, ALS18 and ALS14 exhibited a strong VHL stabilization effect, mirroring an effect previously observed with VHL inhibitors VH032 and VH298.^48^ Among the new dimerizers, ALS14 showed the least stabilization of pVHL30 at higher concentrations, consistent with its weaker cellular VHL engagement and weaker ternary complex association *in vitro*. This is also consistent with our ubiquitination data: although ALS14 promotes the accumulation of monoubiquitinated VHL with UBE2D2, in a cellular context more than one E2 and E3 species could work in tandem to further extend the ubiquitin chains, therefore increasing the likelihood of forming K48-linked tetraubiquitin chains for recognition and degradation by the 26S proteosome. As such, the reduced stabilization mediated by ALS14 could reflect its ability, albeit low, to mediate ubiquitination and degradation in a cellular context.

In addition to VHL, protein levels of Cul2 and its neddylated form NEDD8-Cul2 were assessed. As previously shown,^35^ CM11 effectively reduced Cul2 levels, more prominently at 24 hours than at 4 hours (**Figures 5A**, **5C** and **S14**). In contrast, none of the newly tested VHL-dimerizers reproduced this effect to the same extent, showing only minor reduction in Cul2 protein level after 24 hours treatment (**Figure 5C**).

Regarding downstream effects, the functional consequence of VHL inhibition/degradation was evaluated by monitoring the protein levels of HIF-1α and its hydroxylated form (HIF-1α-OH). The most striking observation was that CC124, ALS18 and ALS14 was mirrored by accumulation of HIF-1α-OH (and total HIF-1α) protein levels, at concentrations of 10 μM or higher, already at 4 h (**Figure S14**) and crucially maintained at 24 h (**Figure 5A**). This suggested that the VHL stabilization induced by the new dimerizers is also inhibitory for VHL activity. While this is consistent with the acute effect of VHL inhibitors such as VH032 and VH298 which induce marked rapid stabilization of HIF-α-OH,^21,49^ it contrasts their longer-term effects which lead to feedback reduction in levels of HIF-1α protein.^48^ As control, no major accumulation in HIF-1α-OH protein level was observed upon CM11-treatment in cells, due to pVHL19 levels remain largely unaffected.^35^

Together, our data support the hypothesis that, despite retaining formation of stable long-lived ternary complexes, the design of more constrained VHL dimerizers abrogated VHL ubiquitination, turning a VHL-degrading homoPROTAC into inhibitory VHL stabilizers.

## Discussion

Protein degraders harness Cullin RING E3 ligases to ubiquitinate and eliminate target proteins.^1,7^ Efficient degradation often arises from stable, long-lived ternary complexes that form at low degrader concentrations,^23–29^ and that position the recruited substrate for productive lysine ubiquitination.^33,34^ PROTAC potency can be enhanced by rigidifying linkers to lock bifunctional compounds into active conformations, a strategy that has yielded potent degraders.^50–53^ Here, we used this strategy to probe the consequences of stabilizing a sub-optimal ternary complex. We solved a cryo-EM structure of the CRL2^VHL^ ligase dimerized by the VHL degrader CM11. To further lock the observed orientation—directed away from the E2∼Ub catalytic site—we designed and synthesized linker-rigidified, sterically constrained VHL homodimerizers. These compounds formed long-lived, compact ternary complexes that were non-productive for ubiquitination, resulting in robust VHL stabilization. Our findings provide a proof-of-concept that protein degraders can be reprogrammed into stabilizers by enforcing non-ubiquitinable ternary complexes. The balance between degradation and stabilization was dictated by linker pattern and length. The head-to-head CM11 symmetric homo-PROTAC was captured in a non-ubiquitinable ternary complex yet, owing to its long, flexible linker, still accesses conformations that enable VHL cross-ubiquitination and degradation. By contrast, side-by-side designs split outcomes: ALS14 induce only partial (non-degradable) ubiquitination, whereas ALS18 and CC124 enforce the most compact geometries and fully abrogate VHL ubiquitination. In cell, ALS18 and CC124 stabilize VHL in an inhibited state, concomitant with increased HIF-1α levels. Thus, VHL dimerization via linker rigidification could be useful where blocking VHL to raise HIF levels and trigger a hypoxic response confers cellular or therapeutic benefits.^21^

Together, this study suggests key principles governing substrate ubiquitinability and degradability (**Figure 6**). In favored systems such as the CRL2^VHL^-MZ1-BRD4^BD2^, a stable, long-lived ternary complex positions lysine optimally for ubiquitination, yielding fast, potent degradation (**Figure 6A**). In such cases, linker rigidification can further enhance PROTAC potency. With the homo-PROTAC CM11, linker flexibility allows the dimerized CRL2^VHL^ to sample multiple conformations; together with cullin flexibility enhanced by neddylation,^39–41^ CM11 can promote ubiquitination on the less favorable VHL substrate (**Figure 6B**). By contrast, rigidifying the linker to trap a non-ubiquitinatable geometry abrogated ubiquitination and reprogrammed a degrader into an inhibitory stabilizer (**Figure 6C**).

**Figure 6.**
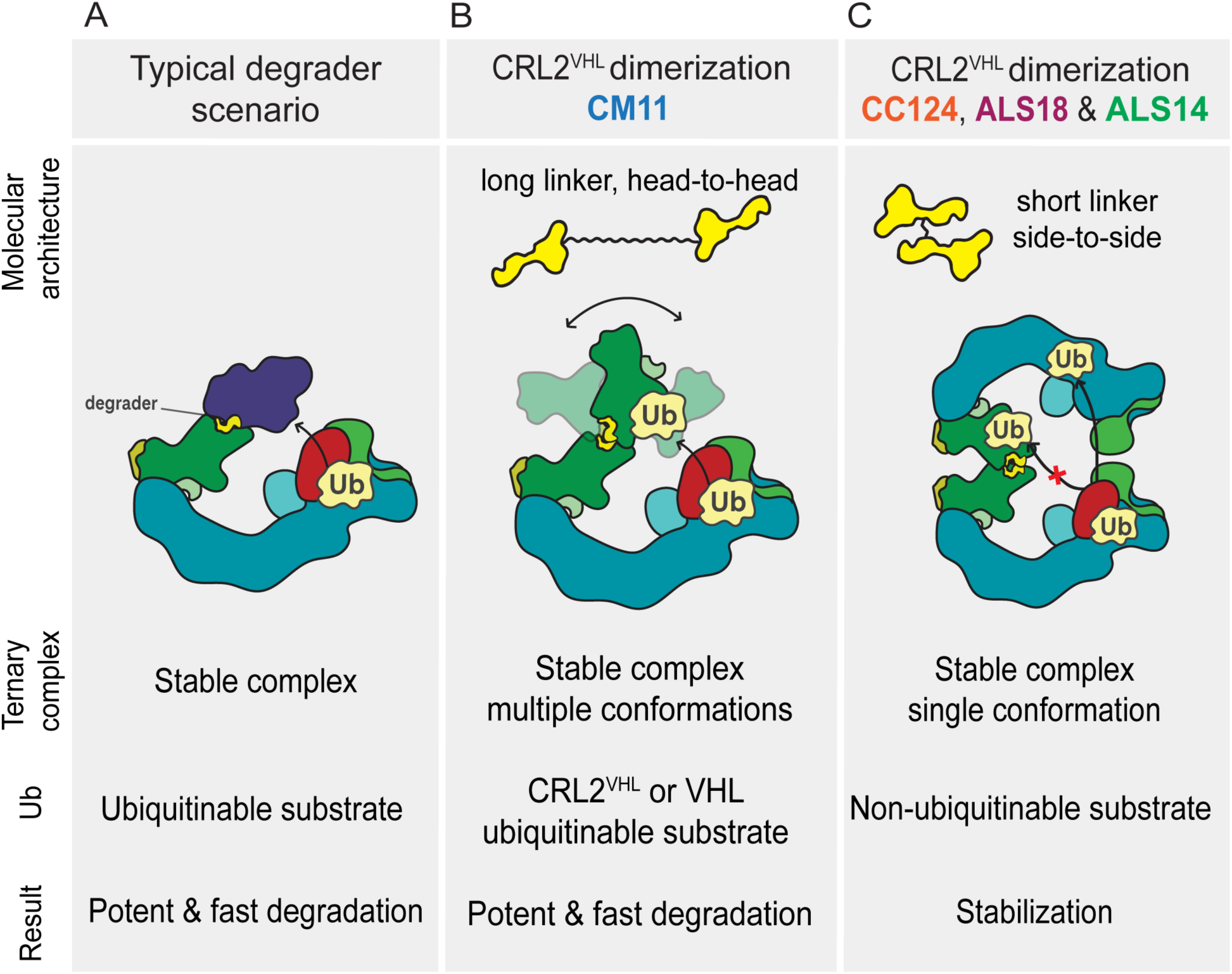
Schematic of key ternary complex scenarios for substrate ubiquitinability.

These insights advance the mechanistic basis of PROTAC-mediated ubiquitinability and degradability, exemplified here by converting degraders into stabilizers when ubiquitinability is occluded by locking the substrate away from the E2∼Ub site. Although we focused on CRL2^VHL^ homodimerizers, the principles should generalize across E3-target pairs. More broadly, tuning ternary complex geometry through linker design can control the functional outcome of induced-proximity modalities, enabling deliberate switching of downstream effects on the recruited target.

## Acknowledgements

We acknowledge the technical and research staff of the Ciulli Laboratories at CeTPD for the set-up and upkeep of protein purification and computational infrastructure. We thank Claudia Diehl for helpful discussions about homoPROTACs; Axel Knebel (University of Dundee) for helpful discussions on protein expression and purification of cullin RING E3 ligases; Xing Liu (Purdue University) for the gift of full-length Cul2-Rbx1 plasmid DNA; Emma Branigan (University of Dundee) for the gift of GACG-ubiquitin plasmid DNA; Mark Nakasone for the gift of NEDD8, APPBP1-UBA3 and UBE2M plasmid DNA; Mark Nakasone and Sarah Chandler (University of Dundee) for the gifts of purified recombinant E1 and E2 for assays; Ramasubramanian Sundaramoorthy for access and support with the Dundee’s School of Life Sciences in-house Cryo-EM Facility; and Mark Norley, Graham Vennall, Robert Arnold, Emily Babcock and Gareth Hughes (Tocris Bio-Techne) for discussions on chemical synthesis.

## Author contributions

C.C. and A.C. conceived the idea, planned the study, and designed compounds. C.C. produced recombinant proteins, performed cryo-EM experiments and solved structures. C.C. A.S. and G.M. designed synthetic routes, with input from H.M. C.C. and A.S. synthesized compounds. C.C., A.S. and G.S. performed assays. C.C. and A.C. co-wrote the manuscript with input from all authors. A.C. acquired research funds and directed the project. All authors have reviewed and approved the final version of the manuscript.

## Data and materials availability

The cryo-EM maps have been deposited to the EMDB under the accession codes EMBD-55480 and EMDB-55558. The atomic model coordinates have been deposited under the accession code PDB-9T32.

