## Supporting Information for "Linker-rigidified VHL homodimerizers convert degraders into stabilizers of non-ubiquitinable ternary complexes"

#### SUPPLEMENTARY FIGURES

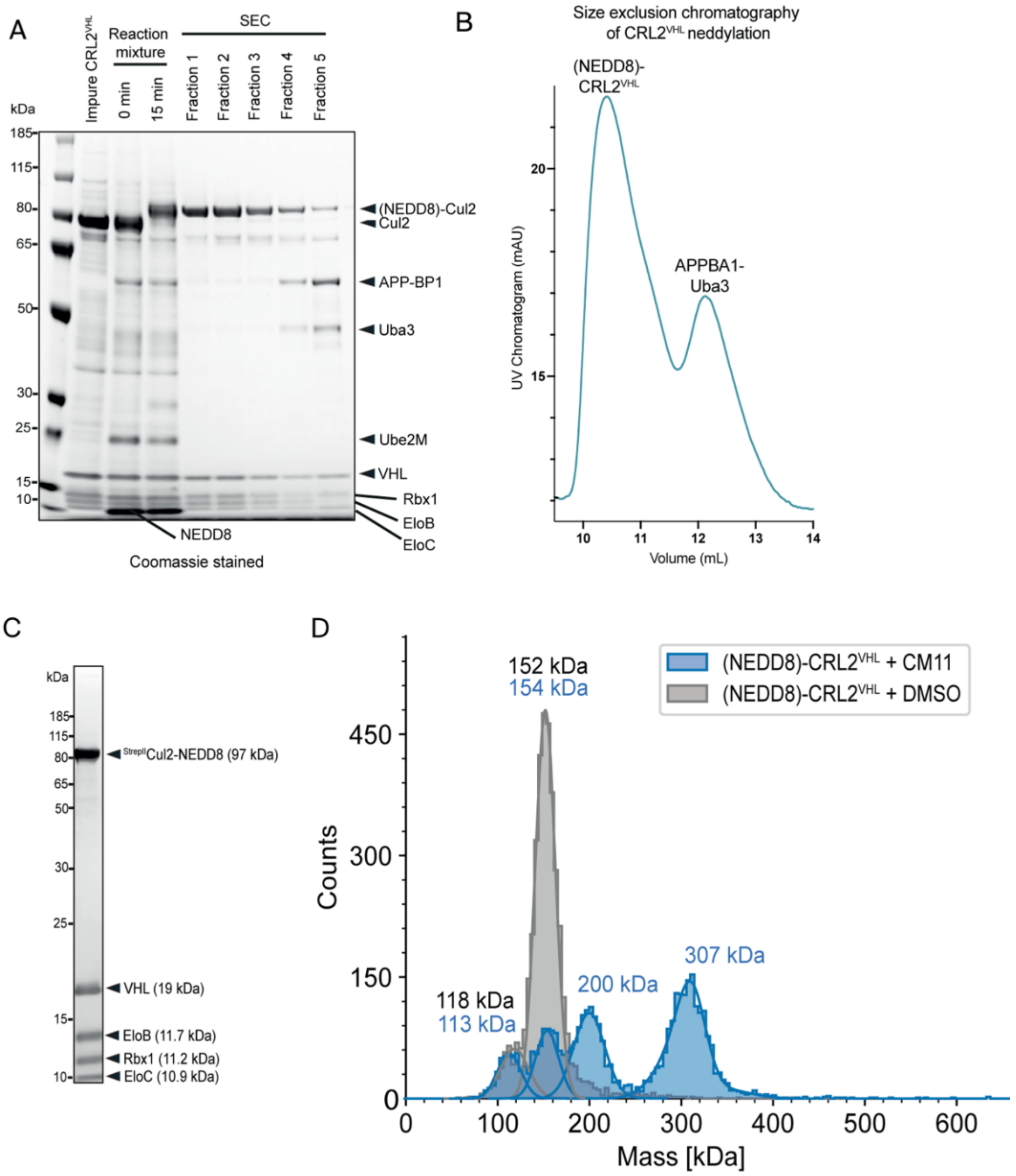

##### Figure S1

**Sample and complex formation for cryo-EM grid preparation**. **A** SDS-PAGE of CRL2$^{VHL}$ neddylation reaction and (NEDD8)-CRL2$^{VHL}$ purification fractions by size exclusion chromatography, adapted from Crowe *et al.*[1] **B** Size exclusion chromatography purification UV trace showing elution of (NEDD8)-CRL2$^{VHL}$ and APPBP1-Uba3 (cropped to elution volumes of 9-14 mL), adapted from Crowe *et al.*[1] **C** SDS-PAGE of the purified (NEDD8)-CRL2$^{VHL}$ used for cryo-EM sample preparation. **D** Mass photometry of (NEDD8)-CRL2$^{VHL}$ alone (green) and of the CM11-dimerised (NEDD8)-CRL2$^{VHL}$ sample applied to cryo-EM grids (blue).

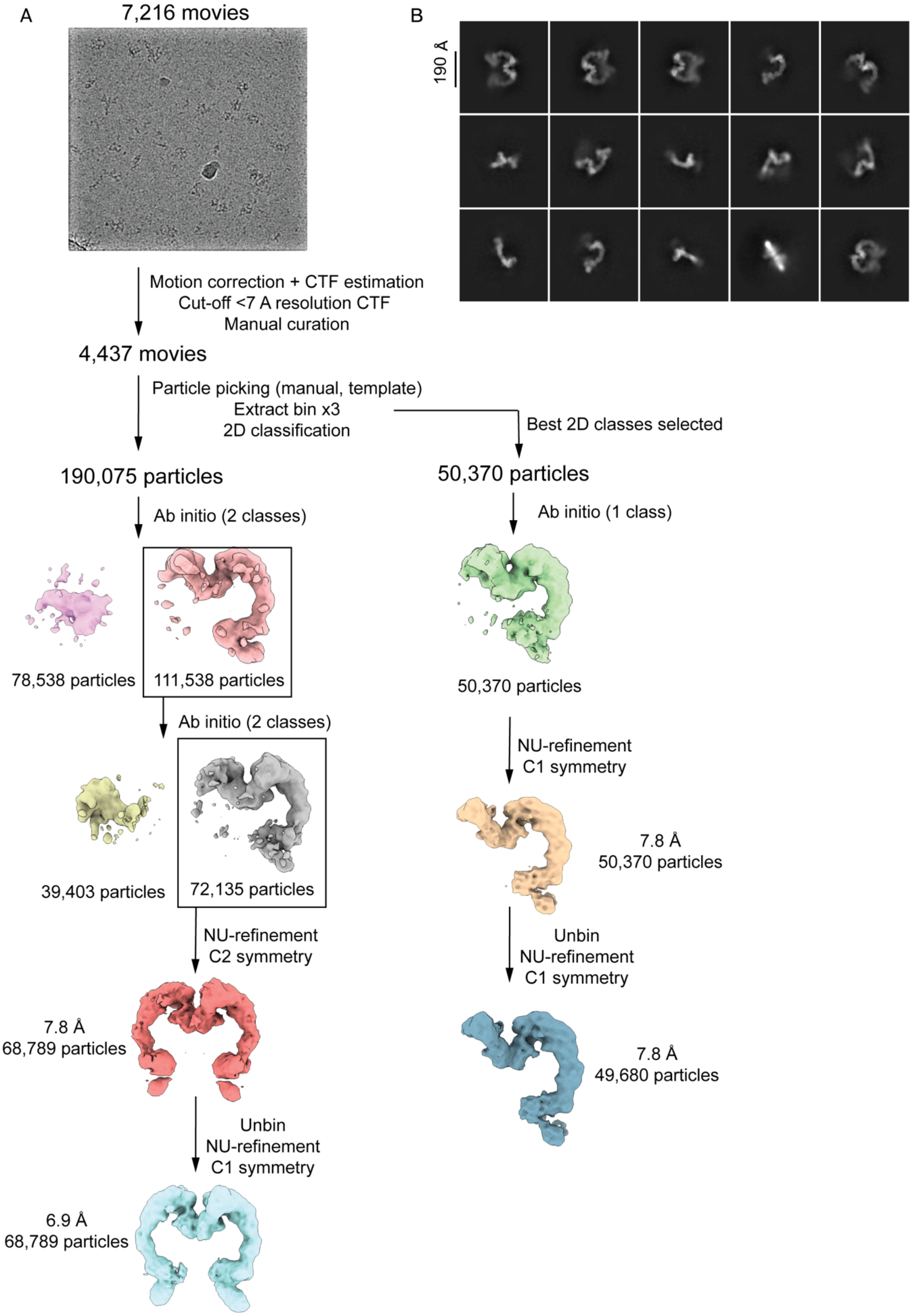

**Figure S2** (continued on next page).

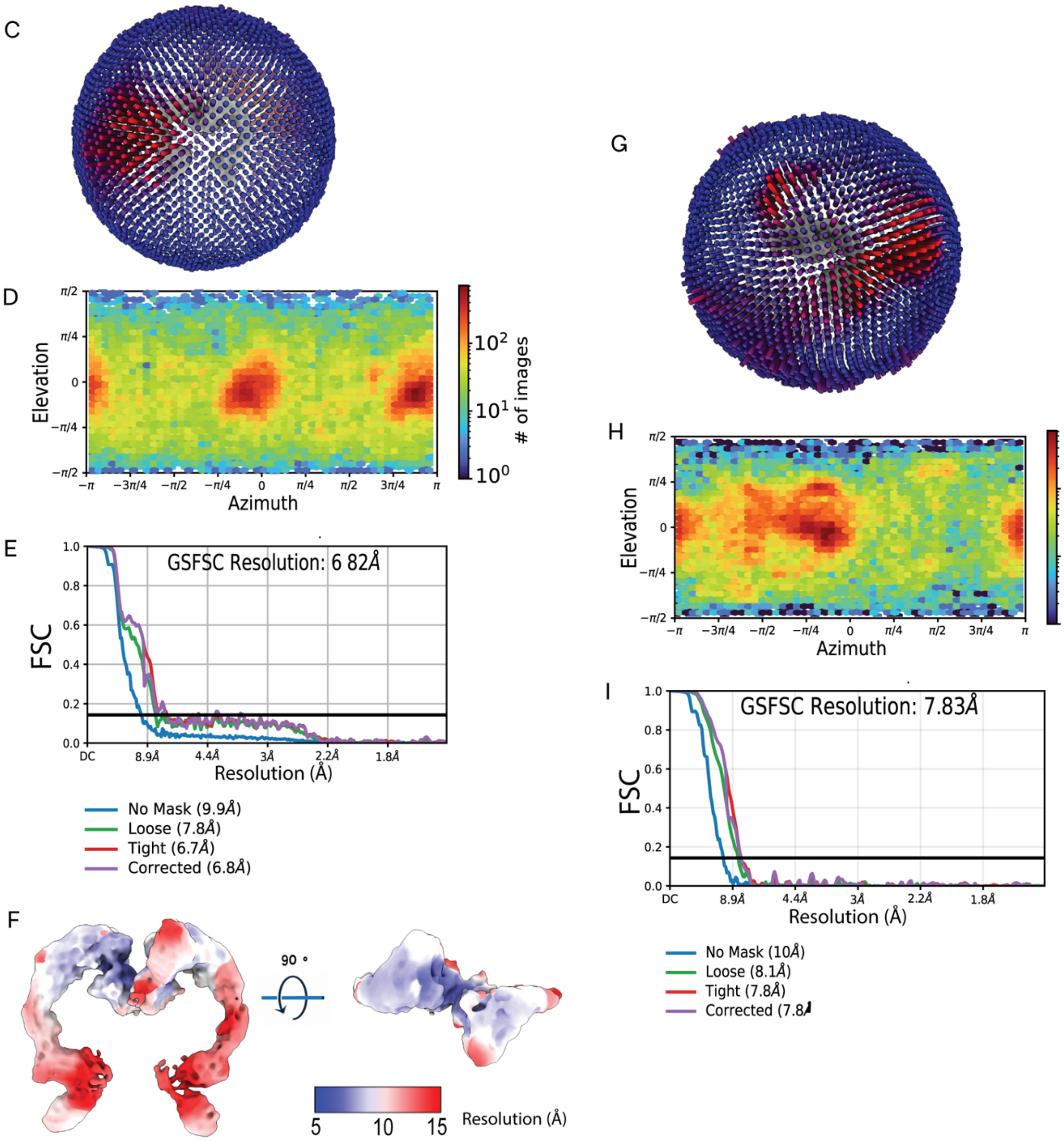

**Figure S2**

**A** Cryo-EM image processing. **B** Representative 2D classes. **C** 3D viewing distribution for the 2D symmetry-masked volume. **D** 2D viewing direction distribution for the 2D symmetry-masked volume. **E** Gold-standard Fourier shell correlation plot at 0.143 for the 2D symmetry-masked volume. **F** Local resolution estimation for the 2D symmetry-masked volume. **G** 3D viewing distribution for the unmasked C1 volume. **H** 2D viewing direction distribution for the unmasked C1 volume. **I** Gold-standard Fourier shell correlation plot at 0.143 for the unmasked C1 volume.

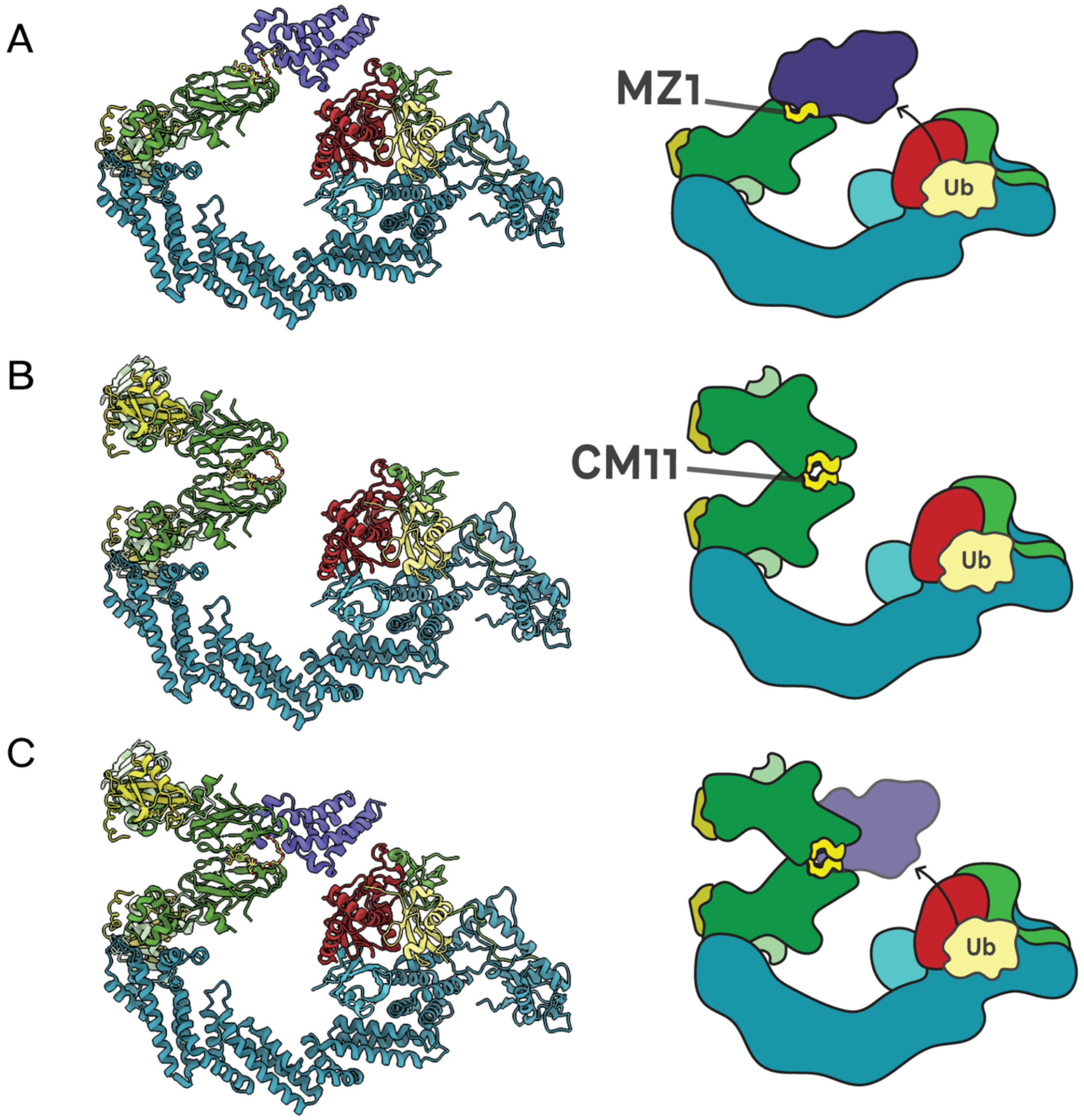

**Figure S3**

**A** Ribbon diagram and corresponding schematic of the cryo-EM BRD4$^{BD2}$-MZ1-(NEDD8)-CRL2$^{VHL}$-UBE2R1-Ub structure PDB: 8RX0[1] showing how MZ1 orients BRD4$^{BD2}$ towards the UBE2R1-Ub catalytic site for ubiquitination. **B** The cryo-EM model in this study aligned with CRL2$^{VHL}$-UBE2R1-Ub from PDB 8RX0, showing how CM11 orients VHL away from the UBE2R1-Ub catalytic site. **C** Superposition of both PDB 8RX0 and the cryo-EM structure in this study. The domains are coloured: VHL (forest green), EloC (light green), EloB (khaki), Cul2 (turquoise), Rbx1 (lime green), BRD4$^{BD2}$ (purple), NEDD8 (cyan), UBE2R1 (red) and ubiquitin (yellow).

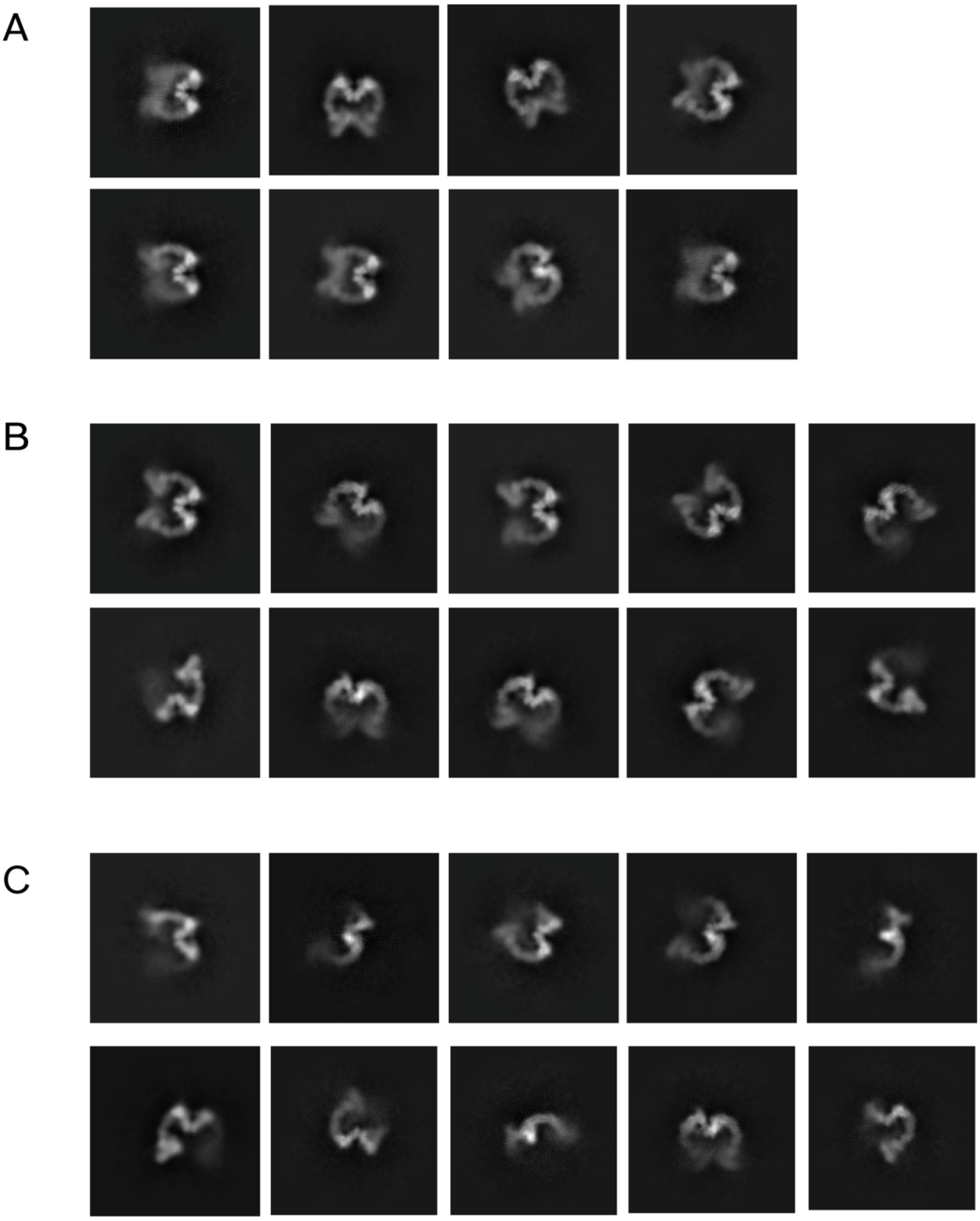

**Figure S4**

**Heterogeneity in the 2D classes from the cryo-EM dataset. A** Selected 2D classes where the cullin C-termini appear close together. **B** Selected 2D classes where the cullin C-termini appear further apart. **C** Selected 2D classes which do not appear similar to classes A or B and suggesting some twisting of one cullin out of the plane.

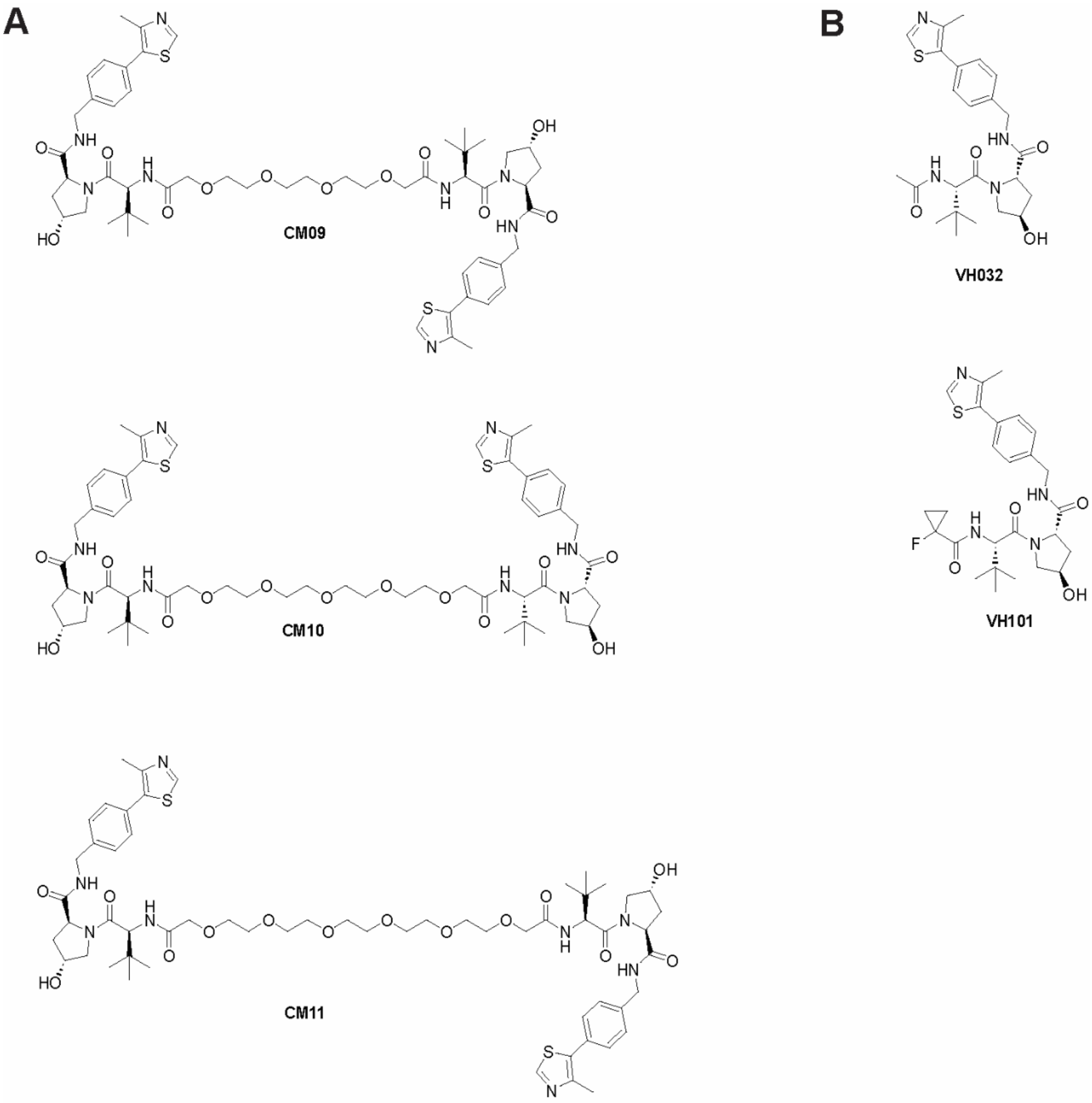

**Figure S5**

**A** CM11 and other previously explored homoPROTAC (CM10 and CM09) endowed with shorter linker lengths **B** VHL ligands used in homoPROTAC design**.** VH032, employed in earlier generations of VHL homoPROTACs, and VH101, used as the ligand scaffold in the design of the VHL dimerizers described in this study.

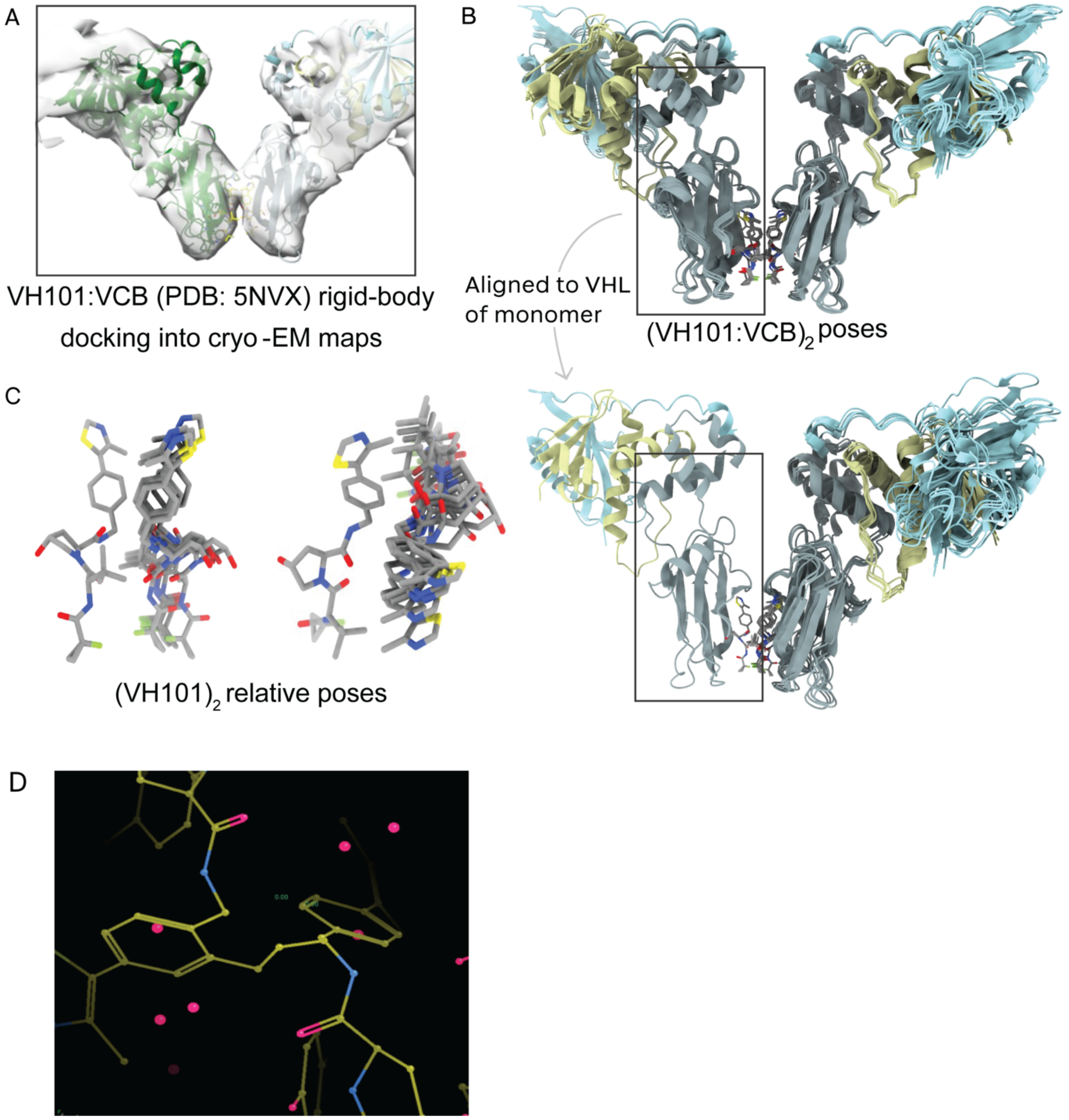

**Figure S6**

**Modelling of the VH101 into cryo-EM maps from this study**. **A** Rigid-body docking of the VHL subunit of the co-crystal structures into 15 independently processed and refined cryo-EM maps. **B** The $(VH101:VCB)_2$ relative poses were aligned onto a single VHL monomer. **C** Alignment to a single monomer allowed us to obtain a distribution of $VH101_2$ poses. **D** Manual modelling revealed an atom-efficient connection to link the two VH101 moieties.

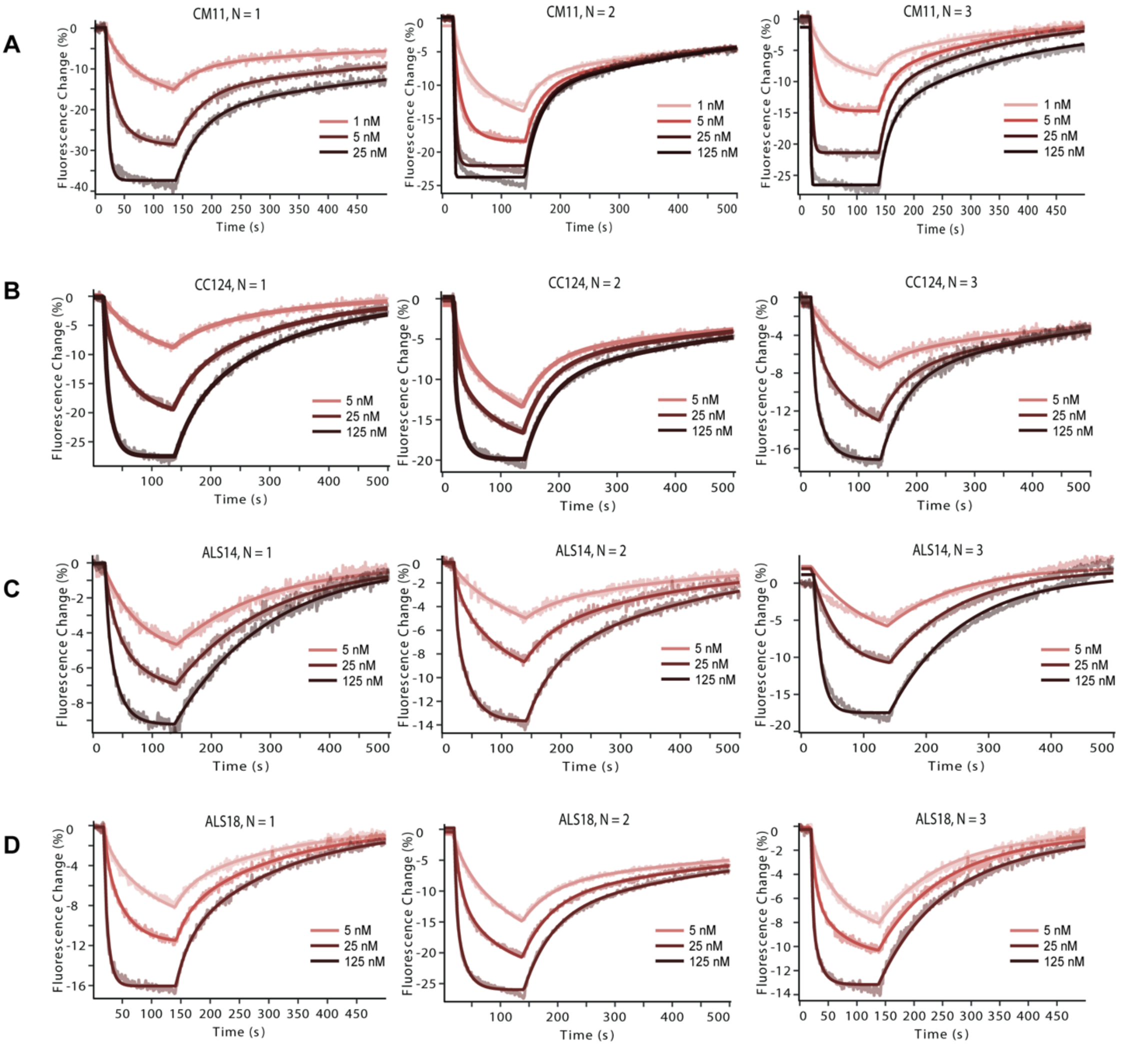

**Figure S7**

Binary binding to VCB measured using the heliX Adapter strands, in N = 3 replicates for **A** CM11; **B** CC124; **C** ALS14; and **D** ALS18.

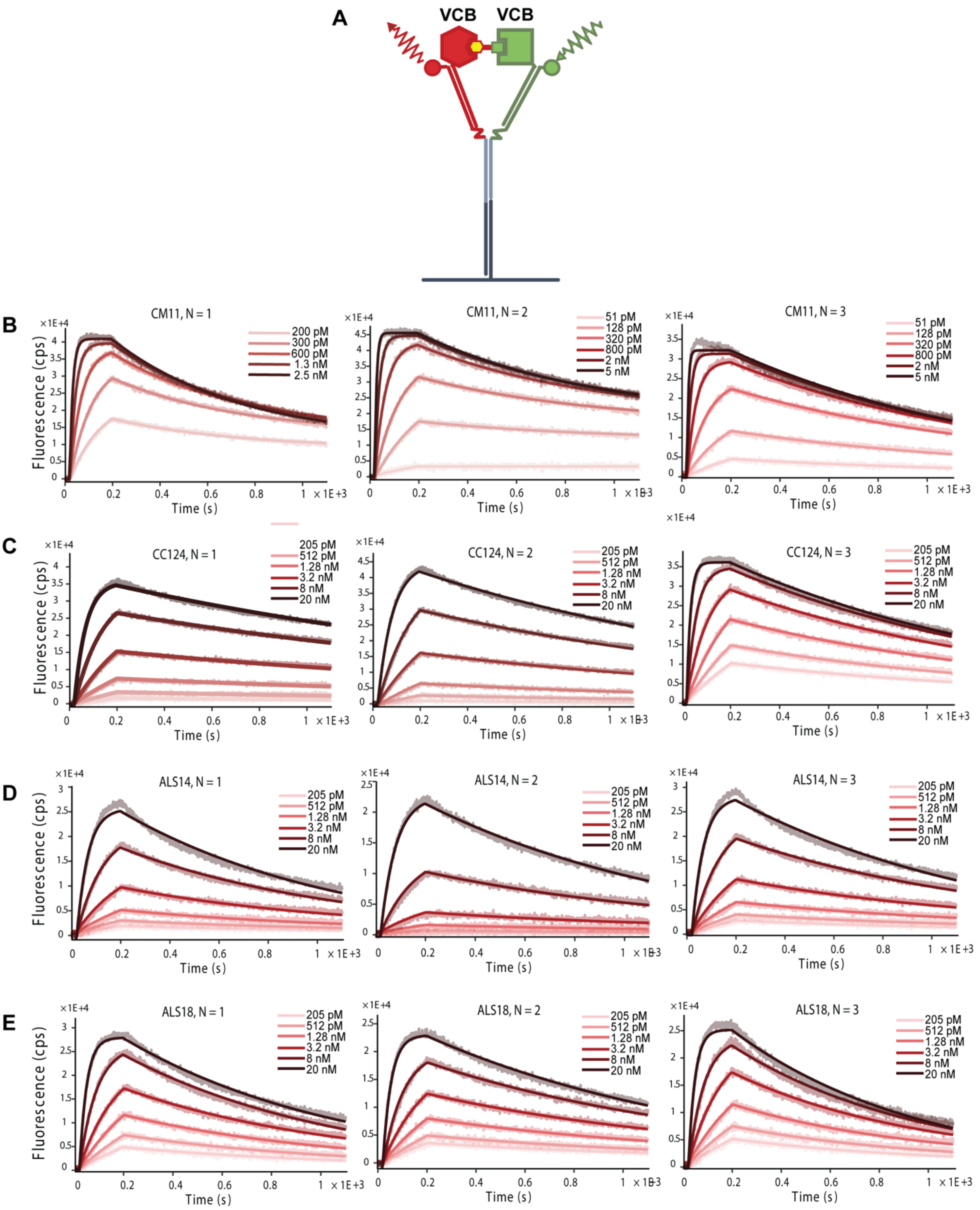

**Figure S8**

**Ternary complex formation between the homoPROTACs in this study and two distinct molecules of VCB. A** Schematic illustrating the assay set-up using the heliX Y-structure. N = 3 replicates are shown for **B** CM11; **C** CC124; **D** ALS14; and **E** ALS18.

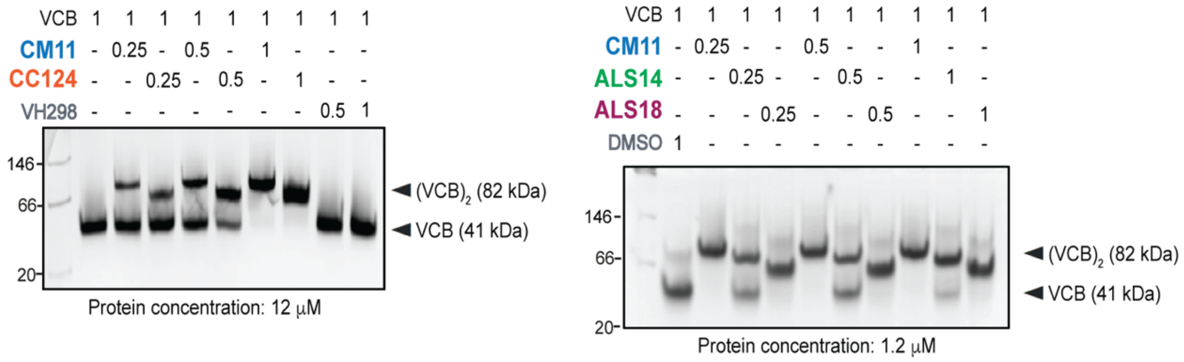

**Figure S9.**

**EMSA showing the dimerization of VCB by CM11, CC124, ALS14 and ALS18, performed at two different protein concentrations (12 μM and 1.2 μM).** A difference in migration was observed between the different compounds with the VCB dimer presenting at slightly different heights in the native gel.

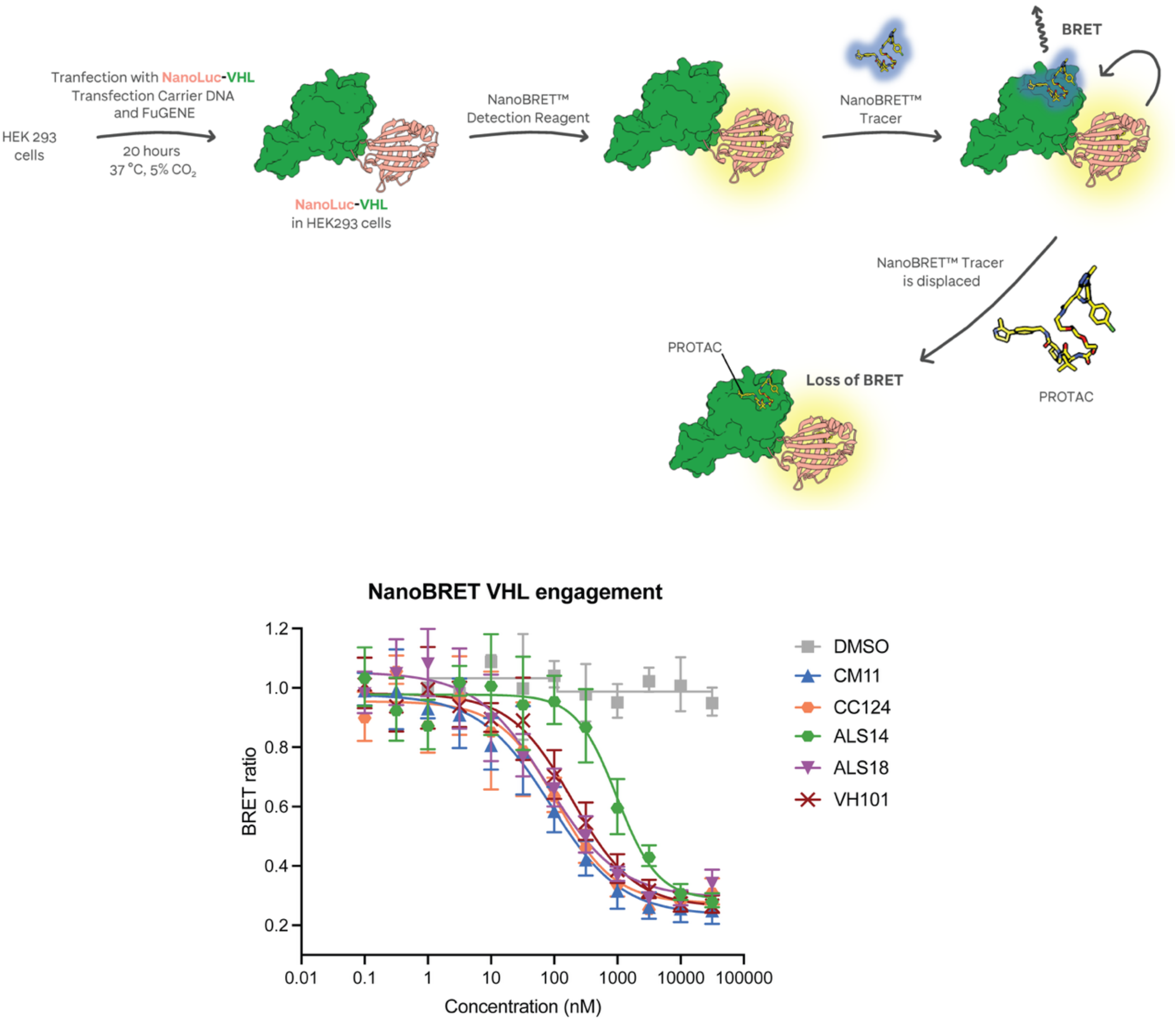

**Figure S10.**

**VHL engagement in digitonin-permeabilized cells measured using a NanoBRET™ system.** HEK293 cells were transfected with DNA encoding for NanoLuc-VHL, permeabilized with digitonin then treated with NanoBRET™ Detection Reagent and NanoBRET™ Tracer to generate a BRET signal. Displacement of the NanoBRET™ Tracer by VH101, CM11, CC124, ALS14 and ALS18 lead to a loss of BRET signal.

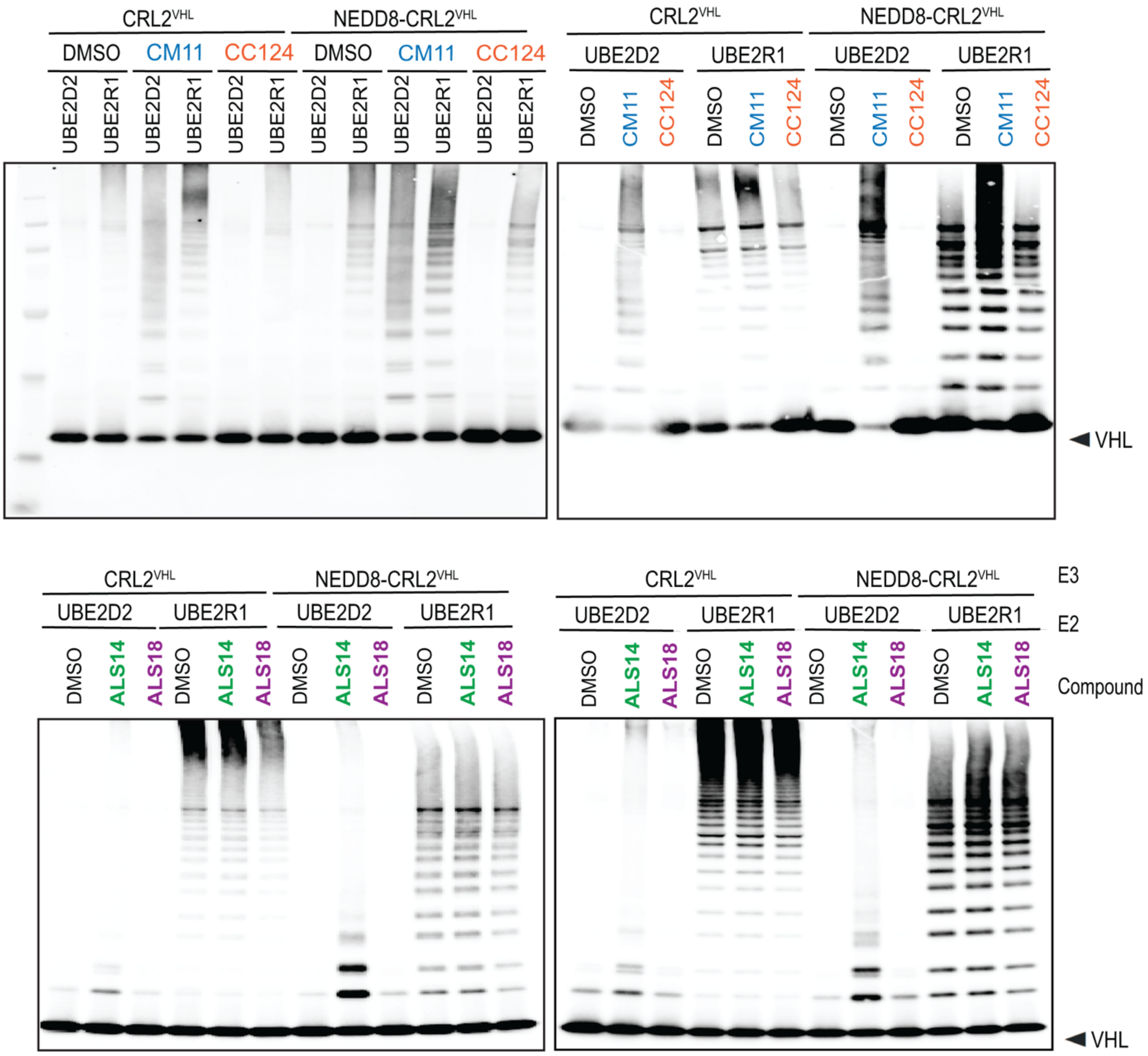

**Figure S11.**

Western blots against VHL for the *in vitro* ubiquitination assays with the homoPROTAC CM11 and the dimerizers CC124, ALS14 and ALS18.

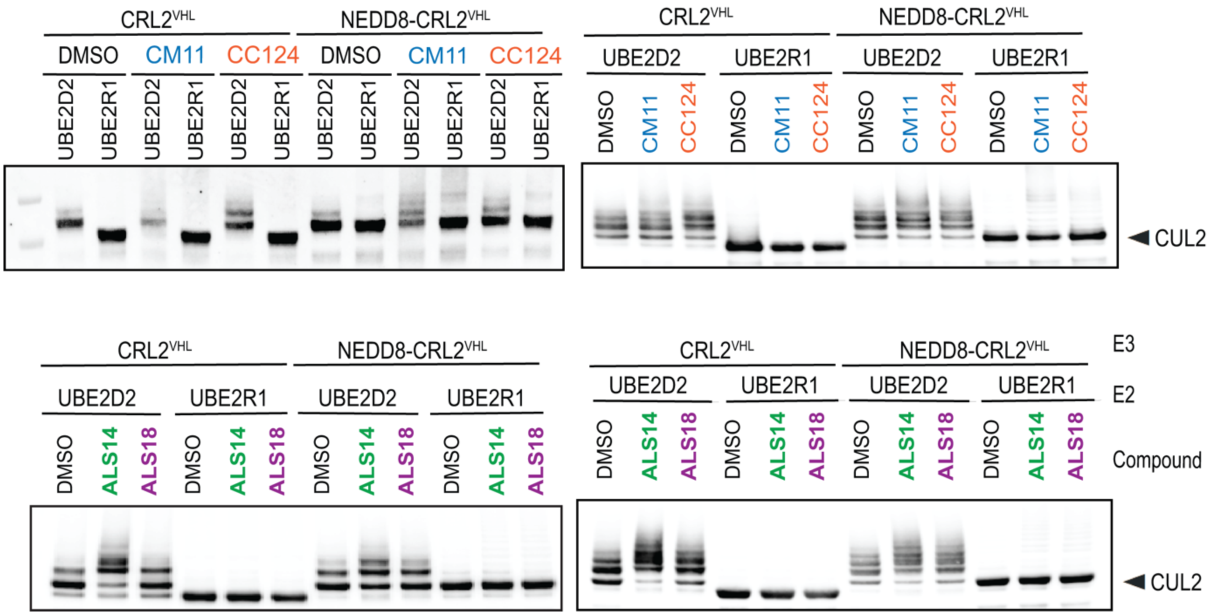

**Figure S12**.
Western blots against Cul2 for the *in vitro* ubiquitination assays with the homoPROTAC CM11 and the dimerizers CC124, ALS14 and ALS18.

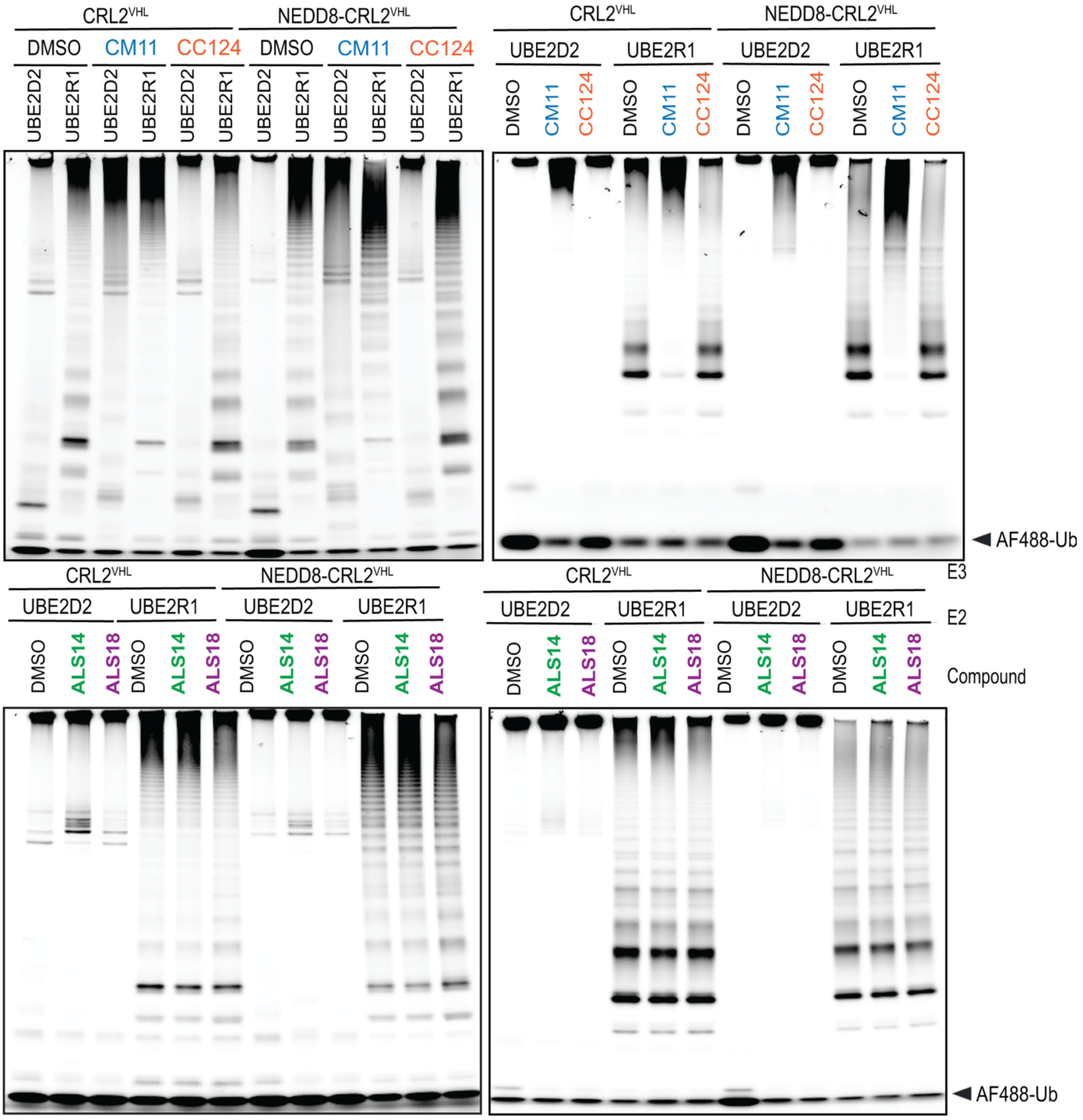

**Figure S13.**

**488 nm imaging of the SDS-PAGE gels for the *in vitro* ubiquitination assays with the homoPROTAC CM11 and the dimerizers CC124, ALS14 and ALS18.** Imaging at this wavelength allows for the detection of the Alexa Fluor 488-labelled ubiquitin species. In all cases, an accumulation of free polyubiquitin chains can be observed throughout the gel and aggregated in the well (top band).

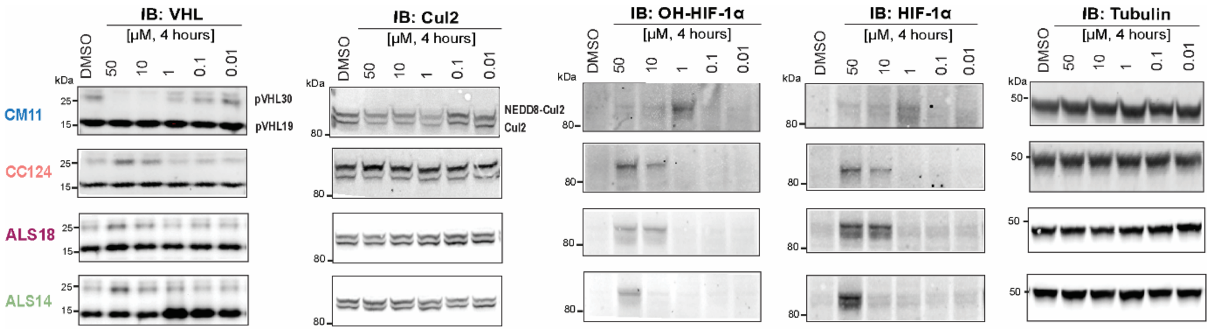

**Figure S14.**

**CC124, ALS14 and ALS18 do not degrade VHL in cells.** Representative immunoblots of dose-dependent effect of compounds CM11, CC124, ALS18 and ALS14 on VHL, Cul2, HIF-1α and OH-HIF-1α in HEK293 cells (representative of $n = 2$ biological replicates, at indicated concentrations, 4 h)

| | The Cullin 2 RING VHL E3 ligase dimerized by the homoPROTAC CM11 *(EMDB-55480, PDB-9T32)* | The Cullin 2 RING VHL E3 ligase dimerized by the homoPROTAC CM11, C1 symmetry *(EMDB-55558)* |
|---|---|---|
| ***Data collection*** | | |
| *Microscope* | Glacios | Glacios |
| *Detector* | Falcon 4i (counting) | Falcon 4i (counting) |
| *Voltage (kV)* | 200 | 200 |
| *Magnification (nominal)* | 190,000x | 190,000x |
| *Number of frames collected* | 640 | 640 |
| *Total electron exposure ($e^-/Å^2$)* | 57 | 57 |
| *Defocus range (μm)* | -1.7 to -3.2 μm | -1.2 to -3.0 μm |
| *Pixel size at detector (Å/pixel)* | 0.74 | 0.74 |
| *Total exposure (s)* | 4.0 | 4.0 |
| *Automation software* | EPU | EPU |
| *Energy filter slit width (eV)* | N/A | N/A |
| *Movies collected (#)* | 7,216 | 7,216 |
| *Movies used (#)* | 4,437 | 4,437 |
| ***Reconstruction*** | | |
| *Image processing package* | CryoSPARC | CryoSPARC |
| *Symmetry imposed* | C1 | C1 |
| *Initial particles (#)* | 111,538 | 50,370 |
| *Final particles (#)* | 68,789 | 49,680 |
| *Map resolution (Å) at FSC 0.143* | 6.9 | 7.8 |
| *B Factor ($Å^2$)* | 306 | 542 |
| *Map resolution range (Å)* | 5.2 to 16.4 | 6.5 to 12.7 |
| *SCF** | 0.901 | 0.960 |
| *cFAR* | 0.02 | 0.20 |
| ***Model composition*** | | |
| *Proteins* | Cul2, Rbx1, EloB, EloC, VHL | |
| *Ligands* | CM11 | |
| ***Model refinement*** | | |
| *Atomic modelling packages* | iSOLDE, Phenix | |
| *Initial model(s) used* | 8RWZ, 4W9H | |
| *Cross-correlation (map/mask)* | 0.55 | |
| *R.m.s deviations from ideal values:* | | |
| *- Bond lengths (Å)* | 0.028 | |
| *- Bond angles (°)* | 2.206 | |
| *Protein residues B Factor(Å)* | 122.66 | |
| *Ligand B Factor (Å)* | 105.30 | |
| ***Validation*** | | |
| *MolProbity score* | 2.33 | |
| *Clashscore (all atoms)* | 10.96 | |
| *Poor rotamers (%)* | 3.41 | |
| *Ramachandran - favoured (%)* | 94.66 | |
| *Ramachandran outliers (%)* | 0.38 | |
| *CaBLAM outliers (%)* | 1.46 | |

#### Table S1.

Cryo-EM data collection, image analysis, atomic modelling, refinement, and validation statistics.

| | VCB-homoPROTAC binary binding | | | | |
|---|---|---|---|---|---|
| *Compound* | $k_{on} \times 10^5$ ($M^{-1}$ $s^{-1}$) | $k_{off} \times 10^{-3}$ ($s^{-1}$) | $K_d$ (nM) | $t_{1/2}$ (s) | *N replicates* |
| *CM11* | 158 ± 44 | 40 ± 14 | 3 ± 2 | 17 | 3 |
| *CC124* | 37 ± 20 | 23 ± 2 | 8 ± 4 | 30 | 3 |
| *ALS14* | 36 ± 23 | 13 ± 10 | 9 ± 5 | 53 | 3 |
| *ALS18* | 41 ± 10 | 25 ± 17 | 6 ± 3 | 28 | 3 |

**Table S2**

**Binary binding results of the compounds used in this study to VHL** (VHL-EloC-EloB construct) using the heliX Adapter strands with measurement in FPS mode.

| | VCB-homoPROTAC-VCB ternary complex formation | | | | |
|---|---|---|---|---|---|
| *Compound* | $k_{on} \times 10^5$ ($M^{-1}$ $s^{-1}$) | $k_{off} \times 10^{-4}$ ($s^{-1}$) | $K_d$ (pM) | $t_{1/2}$ (s) | *N replicates* |
| *CM11* | 316 ± 28 | 8 ± 2 | 25 ± 4 | 886 | 3 |
| *CC124* | 17 ± 13 | 6 ± 2 | 468 ± 246 | 1168 | 3 |
| *ALS14* | 10 ± 1 | 10 ± 1 | 1024 ± 125 | 714 | 3 |
| *ALS18* | 16 ± 2 | 11 ± 2 | 650 ± 72 | 643 | 3 |

**Table S3**

**Ternary complex binding results of the compounds used in this study to VHL** (VHL-EloC-EloB construct) using the heliX Y-structure strands with measurement in FRET mode.

### MATERIALS AND METHODS

#### 1. Chemistry

All reactions were performed under an inert nitrogen atmosphere unless otherwise stated. All solvents and chemical reagents were purchased from commercial sources and used without any further purification.

The reactions were monitored by thin layer chromatography, performed on glass plates pre-coated with silica gel (Analtech, UNIPLATE™ 250 µm / UV254) or using aluminium sheets of silica gel 60, with visualization being achieved using UV light (254 nm) and/or by staining with alkaline potassium permanganate dip. The reactions were also monitored by liquid chromatography-mass spectrometry (LC-MS) using Agilent InfinityLab LC/MSD systems or using an Agilent Technologies 1200 series analytical HPLC connected to an Agilent Technologies 6130 quadrupole LC/MS containing an Agilent diode array detector and a Waters XBridge column (50 mm × 2.1 mm, 3.5 µm particle size).

Purification by flash column chromatography was carried out using Fisher Scientific silica gel 60 Å (35-70 µm), or by using Biotage Selekt, Biotage Isolera, Grace Reveleris or Buchi Pure systems using Biotage Sfär silica prepacked columns (60 µm), or Teledyne Isco Combiflash Rf or Rf200i, using RediSep Rf disposable columns. Purification of products by reversed phase flash chromatography was achieved using a Teledyne Isco Combiflash NextGen 300, using RediSep Rf Gold C18 columns. Final products were purified using a Gilson Preparative HPLC System equipped with a Waters X-Bridge C18 column (100 mm x 19 mm; 5 µm particle size) 212 at a flow rate 25 mL/min. The acidic method constituted elution using a gradient of 5% to 95% acetonitrile in 0.1% formic acid in water, unless differently stated. The basic method constituted elution 5% to 95% acetonitrile in 0.1% ammonia in water.

Nuclear magnetic resonance spectra were recorded on a Bruker Avance III HD spectrometer operating at 400 MHz for $^{1}H$ NMR and 100 MHz for $^{13}C$ NMR, or on Bruker 500 Ultrashield at 500 MHz for $^{1}H$ NMR, or a Bruker Ascend 400 at 400 MHz for $^{1}H$ NMR. Deuterated chloroform, methanol and dimethyl sulfoxide were used as solvents for NMR analysis. $^{1}H$ NMR and $^{13}C$ NMR chemical shifts (δ) are reported in parts per million (ppm) and are referenced to residual protium in solvent and to the carbon resonances of the residual solvent peak respectively. DEPT and correlation spectra were run in conjunction to aid assignment. Coupling constants (*J*) are quoted in Hertz (Hz). Multiplicities are abbreviated as: s, singlet; d, doublet; dd, doublet of doublets; t, triplet; q, quartet; m, multiplet; bs, broad singlet.

##### Scheme S1. Synthesis of CC124

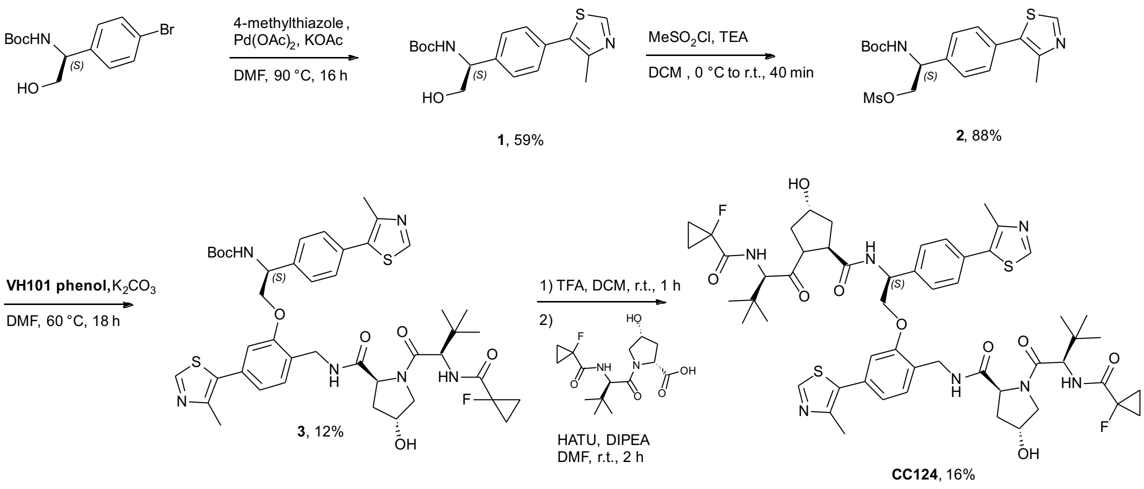

##### Scheme S2. Synthesis of ALS14

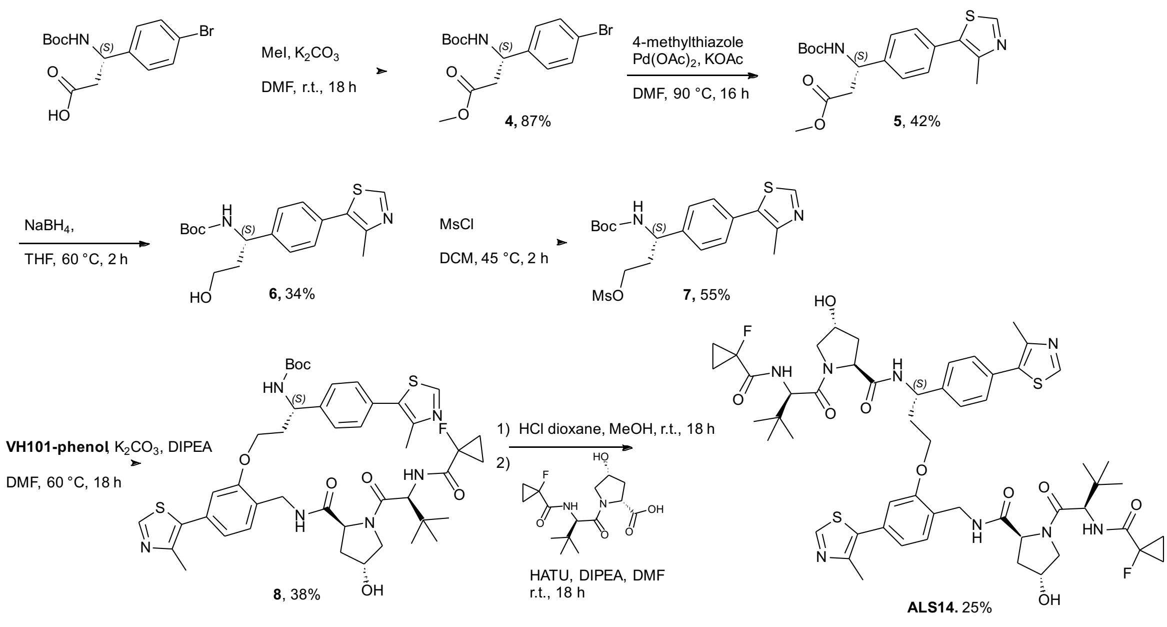

### Scheme S3. Synthesis of ALS18

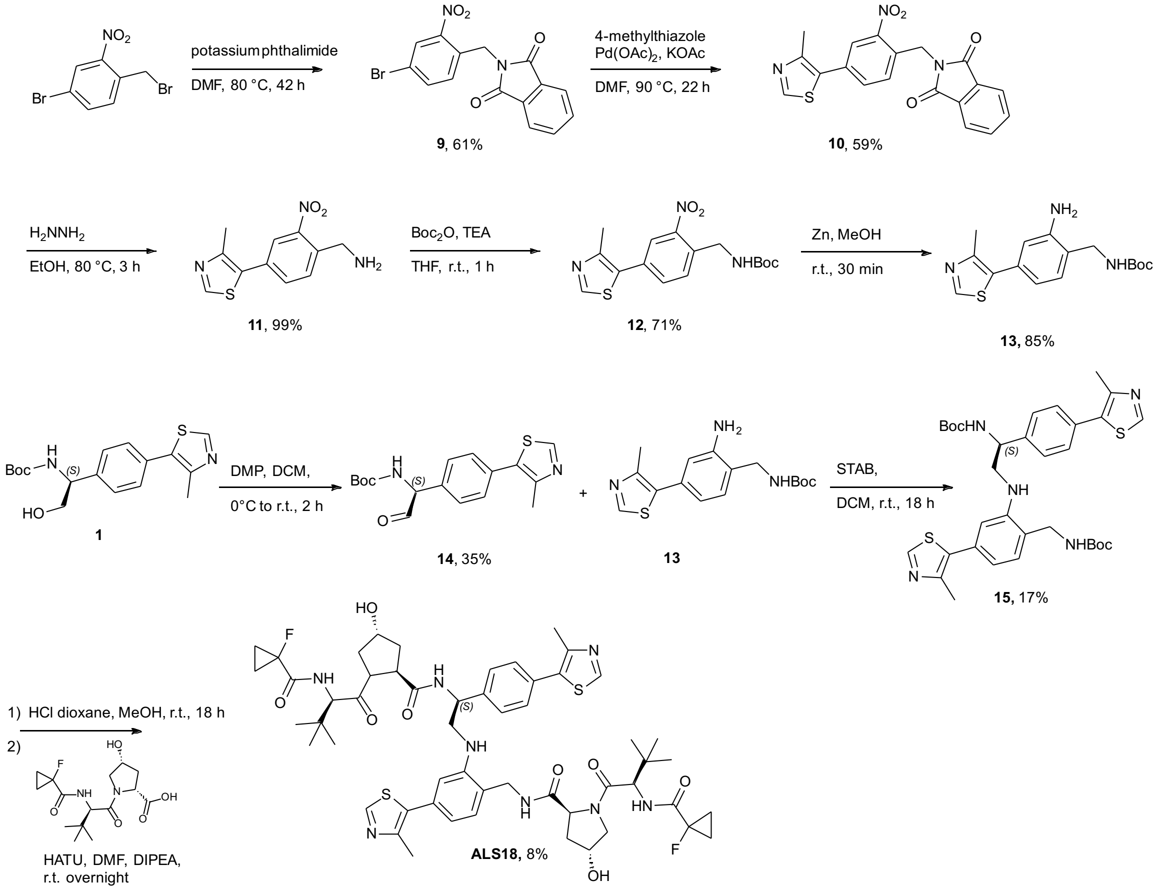

***tert*-Butyl (*S*)-(2-hydroxy-1-(4-(4-methylthiazol-5-yl)phenyl)ethyl)carbamate (1)**

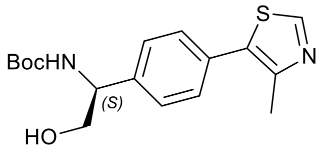

To a stirred solution of *tert*-butyl (*S*)-(1-(4-bromophenyl)-2-hydroxyethyl)carbamate (1.74 g, 5.50 mmol) and potassium acetate (1.08 g, 11.0 mmol) in DMF (15 mL) under argon was added 4-methylthiazole (1.0 mL, 11.0 mmol) and palladium(II) acetate (123 mg, 550 μmol), and the resulting reaction mixture stirred at 90 ºC for 16 hours. After filtration through diatomaceous earth, the filtrate was diluted with water (100 mL) and extracted with ethyl acetate (3 × 25 mL). The combined organic extracts were washed with brine (3 × 250 mL), dried over anhydrous magnesium sulfate and concentrated under reduced pressure to a yellow oil. Purification by flash column chromatography, eluting with 0-60% EtOAc/heptane, afforded the title compound as a yellow solid (1.07 g, 59%). $^{1}$H NMR ($CDCl_3$) $\delta$ 8.69 (s, 1 H), 7.40 (dd, $J$ = 24.7 Hz, $J$ = 8.2 Hz, 4H), 5.32 (d, 1 H), 4.82 (bs, 1 H), 3.94-3.84 (m, 2 H), 2.53 (s, 3 H), 1.45 (s, 9 H). $^{13}$C NMR ($CDCl_3$): $\delta$ 156.1, 150.6, 148.6, 131.7, 131.4, 129.8 (2C), 127.1 (2C), 118.3, 80.3, 66.7, 28.5 (3C), 16.2. *m/z* (ES+): 335.2 $[M+H^+]^+$

**(*S*)-2-((tert-Butoxycarbonyl)amino)-2-(4-(4-methylthiazol-5-yl)phenyl)ethyl methanesulfonate (2)**

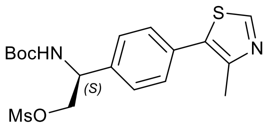

To a stirred solution of **1** (230 mg, 689 μmol) in DCM (1 mL) at 0 ºC methanesulfonyl chloride (66 μL, 826 μmol) was added dropwise followed by triethylamine (143 μL, 1.03 mmol). The resulting reaction mixture was stirred at 0 ºC for 5 minutes, then at room temperature for 40 minutes. The mixture was diluted in water (10 mL) and extracted with DCM (3 × 5 mL). The combined organic extracts were washed with brine (3 × 10 mL), dried over anhydrous magnesium sulfate and concentrated under reduced pressure to a yellow oil (249 mg, 88%). $^{1}$H NMR ($CDCl_3$) $\delta$ 8.71 (s, 1 H), 7.40 (dd, $J$ = 28.4 Hz, $J$ = 8.3 Hz, 4H), 5.20 (bs, 1 H), 5.07 (bs, 1 H), 4.53-4.41 (m, 2 H), 2.97 (s, 3H), 2.54 (s, 3 H), 1.45 (s, 9 H). *m/z* (ES+): 413.2 $[M+H^+]^+$

***tert*-Butyl ((*S*)-2-(2-(((2*S*,4*R*)-1-((*R*)-2-(1-fluorocyclopropane-1-carboxamido)-3,3-dimethylbutanoyl)-4-hydroxypyrrolidine-2-carboxamido)methyl)-5-(4-methylthiazol-5-yl)phenoxy)-1-(4-(4-methylthiazol-5-yl)phenyl)ethyl)carbamate (3)**

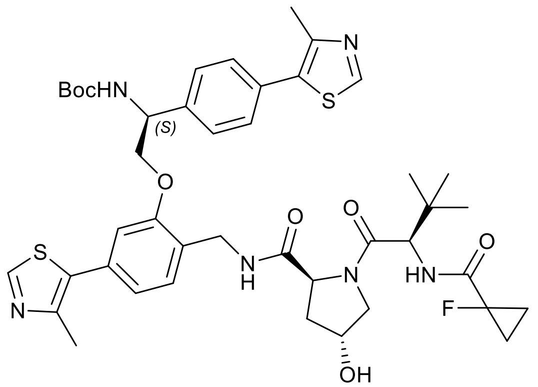

To a stirred solution of **2** (228 mg, 553 μmol) and VH101-phenol (350 mg, 664 μmol) in DMF (1.5 mL) was added potassium carbonate (150 mg, 1.11 mmol). The resulting reaction mixture was stirred at 80 ºC under nitrogen for 18 hours. The reaction mixture was diluted with water (10 mL) and extracted with ethyl acetate (4 × 10 mL). The combined organic extracts were washed with brine (3 × 70 mL), dried over anhydrous magnesium sulfate and concentrated under reduced pressure to a yellow oil. The resulting oil was dissolved in ethyl acetate (20 mL) and washed with 0.1 M NaOH (3 × 20 mL) followed by brine (3 × 100 mL). The organic phase was dried over anhydrous magnesium sulfate and concentrated under reduced pressure to afford the title compound as a yellow solid (55 mg, 12%). $^1$H NMR ($CDCl_3$) $\delta$ 8.67 (s, 1 H), 8.66 (s, 1 H), 7.51 (d, $J$ = 8.2 Hz, 4H), 7.44 (d, $J$ = 8.2 Hz, 4H), 7.29 (d, $J$ = 7.7 Hz, 4H), 6.94 (dd, $J$ = 7.7 Hz, $J$ = 1.4 Hz, 1H), 6.82 (d, $J$ = 1.4 Hz, 1H), 6.07 (bs, 1 H), 5.34 (bs, 1 H), 4.78-4.71 (m, 2 H), 4.54 (bs, 4 H), 4.39-4.35 (m, 3 H), 4.23-4.20 (m, 3 H), 3.93 (d, $J$ = 11.0 Hz, 2H), 3.61 (dd, $J$ = 11.0 Hz, $J$ = 3.5 Hz, 2H), 2.50 (s, 3 H), 2.49 (s, 3 H), 1.43 (s, 9 H), 0.82 (s, 9 H).$^{13}$C NMR ($CDCl_3$) $\delta$ 170.5, 156.39, 150.5, 150.4, 148.6, 148.5, 148.4, 132.4, 131.5, 131.3, 130.3, 129.8, 129.5, 127.1, 126.5, 122.0, 112.2, 118.9, 71.2, 70.1, 58.6, 57.5, 57.3, 56.5, 39.3, 35.3, 28.4, 26.2, 16.1, 16.1, 15.8, 13.8, 13.7. *m/z* (ES+): 749.4 [M-Boc+H$^+$]$^+$

**(2*S*,4*R*)-1-((*S*)-2-(1-Fluorocyclopropane-1-carboxamido)-3,3-dimethylbutanoyl)-*N*-(2-((*S*)-2-((2*S*,4*R*)-1-((*S*)-2-(1-fluorocyclopropane-1-carboxamido)-3,3-dimethylbutanoyl)-4-hydroxypyrrolidine-2-carboxamido)-2-(4-(4-methylthiazol-5-yl)phenyl)ethoxy)-4-(4-methylthiazol-5-yl)benzyl)-4-hydroxypyrrolidine-2-carboxamide (CC124)**

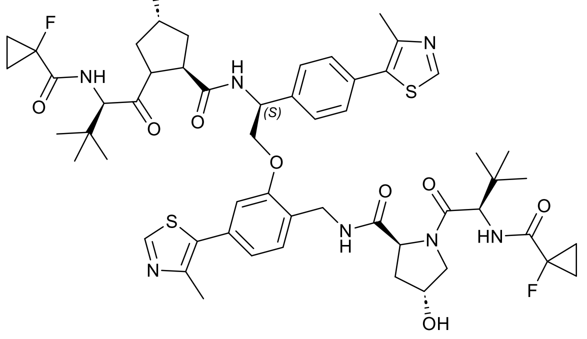

To a stirred solution of **3** (37 mg, 49 μmol) in DCM (0.3 mL) was added TFA (40 μL). The resulting mixture was stirred at ambient temperature under nitrogen for 1 hours. The mixture was treated with DCM/heptane and concentrated under reduced pressure to yield the Boc-deprotected intermediate as a yellow oil. The residue was redissolved in DMF (1 mL) and sequentially treated with (2*R*,4*R*)-1-((*R*)-2-(1-Fluorocyclopropane-1-carboxamido)-3,3-dimethylbutanoyl)-4-hydroxypyrrolidine-2-

carboxylic acid(16.6 mg, 50 μmol), DIPEA (17.5 μL, 100 μmol), HATU (21 mg, 50 μmol) and stirred at ambient temperature for 2 hours. Purification by preparative HPLC, eluting 50-95% 0.1% formic acid in water/acetonitrile, afforded the title product as a white solid (8 mg, 16%). $^{1}$H NMR ($CDCl_3$) $\delta$ 8.92 (d, *J* = 4.8 Hz, 1 H), 8.67 (d, *J* = 2.9 Hz, 1 H), 7.73 (d, *J* = 8.2 Hz, 2 H), 7.54 (dd, *J* = 7.7 Hz, *J* = 5.0 Hz, 1 H), 7.44 (d, *J* = 8.2 Hz, 2 H), 7.33 (d, *J* = 7.7 Hz, 1 H), 7.10 (dd, *J* = 7.7 Hz, *J* = 3.5 Hz, 1 H), 6.91 (dd, *J* = 7.6 Hz, *J* = 1.5 Hz, 1 H), 6.88 (dd, *J* = 9.3 Hz, *J* = 3.5 Hz, 1 H), 6.81 (d, *J* = 1.5 Hz, 1 H), 5.16 (bs, 1 H), 5.05 (dd, *J* = 9.6 Hz, *J* = 6.9 Hz, 1 H), 4.53-4.39 (m, 6 H), 4.34 (d, *J* = 7.6 Hz, 2 H), 3.97 (dd, *J* = 14.0 Hz, *J* = 4.7 Hz, 1 H), 3.82-3.69 (m, 5 H), 3.61 (dd, *J* = 11.0 Hz, *J* = 3.2 Hz, 1 H), 2.54 (s, 3 H), 2.52 (s, 3 H), 2.24-2.17 (m, 1 H), 2.09-2.02 (m, 1 H), 1.99-1.89 (m, 2 H), 1.57 (bs, 2 H), 1.44-1.27 (m, 8 H), 1.04 (s, 9 H), 0.67 (s, 9 H). $^{19}$F NMR ($CDCl_3$) $\delta$ -196.8, -197.4. HRMS (ES+) calculated for $C_{54}H_{67}F_2N_7O_9S_2$ 1061.2868 [M+H$^+$], found 1061.4439

**Methyl (*S*)-3-(4-bromophenyl)-3-((*tert*-butoxycarbonyl)amino)propanoate (4)**

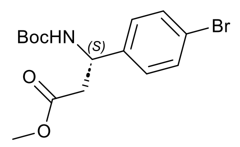

To a stirred suspension of (*S*)-3-(4-bromophenyl)-3-((*tert*-butoxycarbonyl)amino)propanoic acid (39.85 g, 115.8 mmol) and potassium carbonate (29.94 g, 173.4 mmol) in DMF (180 mL) was added methyl iodide (10.7 mL, 173.7 mmol) in a dropwise fashion, and the resulting reaction mixture was stirred at ambient temperature under nitrogen for 18 hours. The reaction mixture was sparged with nitrogen through a methanol/ammonia scrubber, diluted with water (1.8 L) and extracted with ethyl acetate (4 × 250 mL). The combined organic extracts were washed with brine, dried over anhydrous magnesium sulfate and concentrated under reduced pressure. Further drying at 40 °C under vacuum afforded the title compound as a white solid (35.9 g, 87%). $^{1}$H NMR ($CDCl_3$) $\delta$ 7.43 (d, *J* = 8.5 Hz, 2 H), 7.17 (d, *J* = 8.5 Hz, 2 H), 5.54 (bs, 1 H), 5.03 (bs, 1 H), 3.61 (s, 3 H), 2.87-2.75 (m, 2 H), 1.41 (s, 9 H). $^{13}$C NMR ($CDCl_3$) $\delta$ 171.2, 155.0, 140.3, 131.7 (2C), 127.9 (2C), 121.4, 79.9, 51.9, 50.7, 40.4, 28.3 (3C). *m/z* (ES+): 382.1 [M+Na$^+$]$^+$

**Methyl (*S*)-3-((*tert*-butoxycarbonyl)amino)-3-(4-(4-methylthiazol-5-yl)phenyl)propanoate (5)**

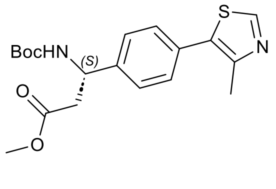

To a stirred solution of **4** (2.51 g, 7.0 mmol) and potassium acetate (1.36 g, 13.9 mmol) in DMF (20 mL) was added4-methylthiazole (1.3 mL, 14.3 mmol) and palladium(II) acetate (701 mg, 3.1 mmol) and the resulting reaction mixture was stirred at 90 ºC under argon for 16 hours.

After filtration through diatomaceous earth, the filtrate was treated with water (200 mL) and extracted with ethyl acetate (4 × 30 mL). The combined organic extracts were washed with brine (3 × 400 mL), dried over anhydrous magnesium sulfate and concentrated under reduced pressure to a yellow oil. Purification by flash column chromatography, eluting with 0-40% EtOAc/heptane, afforded the title compound as a yellow solid (1.12 g, 42%).$^{1}$H NMR ($CDCl_3$) *δ* 8.65 (s, 1H), 7.39 (d, *J* = 8.4 Hz, 2 H), 7.34 (d, *J* = 8.4 Hz, 2 H), 5.61 (bs, 1 H), 5.12 (bs, 1 H), 3.62 (s, 3 H), 2.91-2.80 (m, 2 H), 1.41 (s, 9 H). $^{13}$C NMR ($CDCl_3$): *δ* 171.4, 155.1, 150.4, 146.6, 141.2, 131.5, 131.2, 129.6, 126.6, 79.9, 51.9, 50.9, 40.6, 28.4, 16.1. *m/z* (ES+): 377.1 $[M+H^+]^+$

***tert*-Butyl (*S*)-(3-hydroxy-1-(4-(4-methylthiazol-5-yl)phenyl)propyl)carbamate (6)**

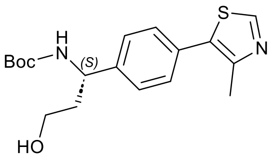

To a stirred solution of **5** (600 mg, 1.59 mmol) in THF (3 mL) was added sodium borohydride (132 mg, 3.50 mmol) portion-wise under an inert atmosphere, followed by MeOH (0.3 mL). The reaction mixture was heated to 60 °C for 2 hours. Upon completion, the reaction was quenched with saturated aqueous ammonium chloride solution and extracted with ethyl acetate (2 × 30 mL). The combined organic extracts were washed with brine (3 × 400 mL), dried over anhydrous sodium sulfate and concentrated under reduced pressure. Purification by flash column chromatography, eluting with 0-5% MeOH/DCM, afforded the title compound as a pale-yellow solid (186 mg, 34%).$^{1}$H NMR (500 MHz, $CDCl_3$) *δ* 8.67 (s, 1H), 7.40 (d, *J* = 8.4 Hz, 3H), 7.35 (d, *J* = 8.1 Hz, 2H), 5.28 – 5.21 (m, 1H), 4.96 – 4.92 (m, 1H), 3.77 – 3.68 (m, 2H), 2.51 (s, 3H), 2.16 – 2.04 (m, 1H), 1.90 – 1.81 (m, 1H), 1.44 (s, 9 H). $^{13}$C NMR (126 MHz, $CDCl_3$) *δ* 156.4, 150.5, 148.6, 142.1, 131.7, 131.1, 130.0, 129.7, 126.9, 126.7, 126.6, 80.2, 59.1, 53.5, 51.7, 39.3, 28.5, 16.2. *m/z* (ES+): 349.0 $[M+H^+]^+$

***(S)*-3-((*tert*-Butoxycarbonyl)amino)-3-(4-(4-methylthiazol-5-yl)phenyl)propyl methanesulfonate (7)**

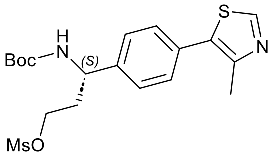

To a stirred solution of **6** (186 mg, 530 µmol) in anhydrous DCM (3 mL) was added methanesulfonyl chloride (91 mg, 1.5 mmol) and DIPEA (279 µL, 1.60 mmol). The sealed vial was heated to 45 °C and stirred for 2 hours under nitrogen. The reaction mixture was poured into water and citric acid (5%), and extracted with DCM (3 × 10 mL). The combined organic extracts were washed with brine, dried over anhydrous sodium sulfate and concentrated under reduced pressure. Purification by flash column chromatography, eluting with 0-100% EtOAc/heptane afforded the title compound (126 mg, 55%). $^1$H NMR (500 MHz, $CDCl_3$) $\delta$ 8.68 (s, 1H), 7.43 (d, $J$ = 8.3 Hz, 2H), 7.34 (d, $J$ = 8.3 Hz, 2H), 4.89 (s, 1H), 4.36 – 4.19 (m, 2H), 3.03 (s, 3H), 2.53 (s, 3H), 2.24 (q, $J$ = 6.7 Hz, 2H), 1.42 (s, 9H). $^{13}$C NMR (126 MHz, $CDCl_3$) $\delta$ 155.3, 150.5, 150.5, 148.8, 131.6, 131.5, 129.9, 126.8, 126.8, 80.2, 66.8, 51.4, 41.4, 39.5, 37.6, 36.0, 28.5, 22.2, 16.2. *m/z* (ES+): 427.0 $[M+H^+]^+$

***tert*-Butyl ((*S*)-3-(2-(((2*S*,4*R*)-1-((*R*)-2-(1-fluorocyclopropane-1-carboxamido)-3,3-dimethylbutanoyl)-4-hydroxypyrrolidine-2-carboxamido)methyl)-5-(4-methylthiazol-5-yl)phenoxy)-1-(4-(4-methylthiazol-5-yl)phenyl)propyl)carbamate (8)**

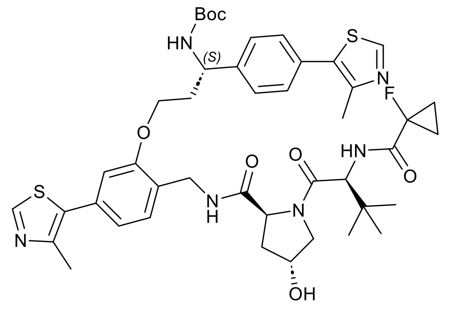

To a stirred solution of VH101-phenol (19 mg, 30 μmol) in anhydrous DMF (1 mL) was added cesium carbonate (11 mg, 30 μmol)and the reaction was stirred for 10 min, followed by the addition of a solution of **7** (15 mg, 30 μmol) in DMF (0.5 mL). The reaction was heated at 60 °C overnight. The reaction mixture was diluted with ethyl acetate (5 mL) and washed with water and brine (3 × 10 mL). The organic layer was washed with brine, dried over anhydrous sodium sulfate and concentrated under reduced pressure. Purification by flash column chromatography, eluting with 0-5% MeOH/DCM afforded the title compound as white powder (10 mg, 38%). $^1$H NMR (500 MHz, $CDCl_3$) $\delta$ 8.68 (s, 1H), 8.67 (s, 1H), 7.43 (s, 4H), 7.35 (d, $J$ = 7.7 Hz, 2H), 6.96 (dd, $J$ = 7.7, 1.6 Hz, 1H), 6.84 (d, $J$ = 1.6 Hz, 1H), 5.73 (s, 1H), 5.03 (s, 1H), 4.79 (s, 1H), 4.54 (s, 1H), 4.52 – 4.42 (m, 3H), 4.03 (d, $J$ = 11.3 Hz, 1H), 3.64 (dd, $J$ = 11.3, 1H), 3.49 (s, 2H), 2.51 (s, 3H), 2.53 (s, 3H), 2.45 – 2.30 (m, 4H), 2.09 (t, $J$ = 10.7 Hz, 1H), 1.37 (s, 9H), 1.30 – 1.27 (m, 4H), 0.90 (s, 9H). $^{13}$C NMR (126 MHz, $CDCl_3$) $\delta$ 170.9, 169.5, 150.3 (2C), 148.5, 126.7 (2C), 121.7, 119.9 (2C), 112.0, 77.3, 77.2, 77.0, 76.8, 60.4, 58.3, 57.6, 53.4, 35.9, 35.5, 35.0, 31.9, 29.0, 29.0, 28.4, 26.3, 25.0, 24.9, 22.7, 21.1, 16.1, 14.2, 14.1. *m/z* (ES+): 864.3 $[M+H^+]^+$

**(2*S*,4*R*)-1-((*R*)-2-(1-Fluorocyclopropane-1-carboxamido)-3,3-dimethylbutanoyl)-*N*-(2-((*S*)-3-((2*S*,4*R*)-1-((*R*)-2-(1-fluorocyclopropane-1-carboxamido)-3,3-dimethylbutanoyl)-4-hydroxypyrrolidine-2-carboxamido)-3-(4-(4-methylthiazol-5-yl)phenyl)propoxy)-4-(4-methylthiazol-5-yl)benzyl)-4-hydroxypyrrolidine-2-carboxamide (ALS14)**

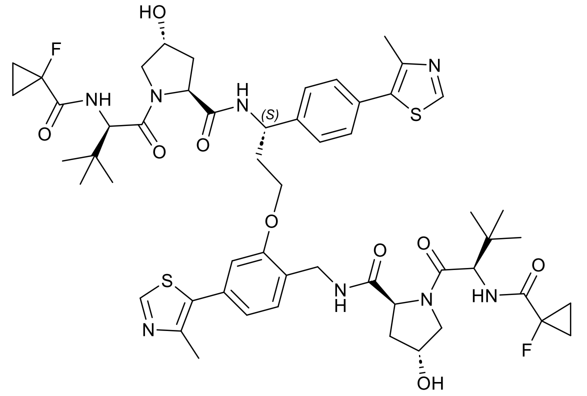

To a stirred solution of **8** (10 mg, 10 µmol) in MeOH (0.5 mL) was added hydrogen chloride (0.5 mL, 4M solution in 1,4-dioxane). The resulting mixture was stirred at room temperature for 18 hours before being concentrated under reduced pressure to yield the Boc-deprotected intermediate as a yellow oil. The residue was redissolved in DMF (0.5 mL) and sequentially treated with (2*R*,4*R*)-1-[(2*R*)-2-[(1-fluorocyclopropanecarbonyl) amino]-3,3-dimethyl-butanoyl]-4-hydroxy-pyrrolidine-2-carboxylic acid (4.5 mg, 0.01 mmol) treated with DIPEA (6.4 µL, 0.03 mmol), HATU (3.8 mg, 0.01 mmol) and stirred at ambient temperature overnight. The reaction mixture was diluted with ethyl acetate c and washed with brine (3 × 10 mL). The organic layer was dried over anhydrous sodium sulfate and concentrated under reduced pressure. Purification by preparative HPLC, eluting with 5-95% 0.1% formic acid in water/acetonitrile, afforded the title compound as a white solid (2.77 mg, 25 %). $^1$H NMR (500 MHz, $CDCl_3$) $\delta$ 8.69 (s, 1H), 8.67 (s, 1H), 8.03 (d, *J* = 9.7 Hz, 1H), 7.77 (s, 1H), 7.49 (d, *J* = 8.0 Hz, 2H), 7.41 (d, *J* = 8.0 Hz, 2H), 7.33 (d, *J* = 7.7 Hz, 1H), 7.04 (dd, *J* = 8.0, 1.7 Hz, 1H), 6.95 (d, *J* = 1.7 Hz, 1H), 6.89 (s, 1H), 6.68 – 6.62 (m, 1H), 5.35 (td, *J* = 10.0, 4.0 Hz, 1H), 4.62 (dd, *J* = 13.0, 3.3 Hz, 1H), 4.54 (dd, *J* = 10.0, 6.6 Hz, 1H), 4.46 (dd, *J* = 10.5, 7.0 Hz, 1H), 4.39 (q, *J* = 10.5 Hz, 4H), 4.32 (d, *J* = 7.1 Hz, 1H), 4.21 (d, *J* = 5.3 Hz, 1H), 4.18 (q, *J* = 5.5 Hz, 1H), 3.94 (d, *J* = 11.1 Hz, 1H), 3.80 (d, *J* = 11.2 Hz, 1H), 3.76 (dd, *J* = 11.2, 3.6 Hz, 1H), 3.67 (dd, *J* = 11.1, 3.7 Hz, 1H), 3.58 (d, *J* = 19.5 Hz, 2H), 2.53 (d, *J* = 10.4 Hz, 6H), 2.50 – 2.38 (m, 1H), 2.18 – 2.11 (m, 1H), 2.05 – 1.96 (m, 2H), 1.93 (dd, *J* = 13.1, 6.9 Hz, 1H), 1.85 (dd, *J* = 13.0, 6.6 Hz, 1H), 1.36 – 1.26 (m, 8H), 1.10 (s, 9H), 0.87 (s, 9H). HRMS (ES+) calculated for $C_{54}H_{68}F_2N_8O_9S_2$ 1075.3018 $[M+H^+]^+$, found 1075.4601

**2-(4-Bromo-2-nitrobenzyl)isoindoline-1,3-dione (9)**

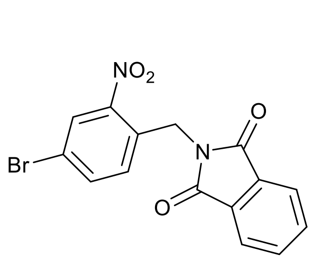

To a stirred solution of 4-bromo-1-(bromomethyl)-2-nitrobenzene (4.98 g, 17.2 mmol) in DMF (19 mL) was added phthalimide potassium salt (3.14 g, 17.2 mmol). The resulting reaction mixture was stirred at 80 ºC for 42 hours. A further portion of phthalimide potassium salt (0.75 g, 4.1 mmol) and DMF (9 mL) was added and the resulting mixture was stirred for 3 hours. The mixture was diluted with water (100 mL) and extracted with ethyl acetate (5 × 30 mL). The combined organic extracts were washed with brine (2 × 200 mL), dried over anhydrous magnesium sulfate and concentrated under reduced pressure to a yellow solid. Trituration with ethyl acetate afforded the title compound as a pale yellow solid (3.76 g, 61%). $^1$H NMR (DMSO-$d_6$) $\delta$ 8.31 (d, $J$ = 2.1 Hz, 1 H), 7.94-7.86 (m, 5 H), 7.45 (d, $J$ = 8.4 Hz, 1 H), 5.05 (s, 2 H). $^{13}$C NMR ($CDCl_3$) $\delta$ 161.1, 148.9, 137.1, 135.1, 132.1 (2C), 131.4, 131.0, 127.9, 123.8 (2C), 121.1. *m/z* (ES+): 401 $[M+K^+]^+$

**2-(4-(4-Methylthiazol-5-yl)-2-nitrobenzyl)isoindoline-1,3-dione (10)**

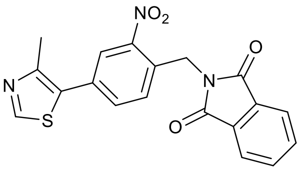

To a stirred solution of **9** (3.76 g, 10.4 mmol) and potassium acetate (2.04 g, 20.8 mmol) in DMF (30 mL) under argon atmosphere were added 4-methylthiazole (1.9 mL, 20.8 mmol) and palladium(II) acetate (233 mg, 1.04 mmol), and the resulting reaction mixture was stirred at 90 ºC for 22 hours. After filtration through diatomaceous earth, the filtrate was treated with water (50 mL) and extracted with ethyl acetate (5 × 50 mL). The combined organic extracts were washed with brine (2 × 500 mL), dried over anhydrous magnesium sulfate and concentrated under reduced pressure to a yellow oil. Purification by flash column chromatography, eluting with 0-100% EtOAc/heptane, afforded the title compound as a yellow solid (2.31 g, 59%). $^1$H NMR (DMSO-$d_6$) $\delta$ 9.03 (s, 1H), 8.10 (d, $J$ = 1.9 Hz, 1 H), 7.88-7.81 (m, 4 H), 7.71 (dd, $J$ = 8.1, $J$ = 1.9 Hz, 1 H), 7.49 (d, $J$ = 8.1, 1 H), 5.07 (s, 2 H), 2.41 (s, 3 H). $^{13}$C NMR ($CDCl_3$) $\delta$ 168.2, 153.4, 150.1, 148.5, 135.1, 134.5, 132.6, 132.1 (2C), 130.9, 130.0, 129.0, 125.2 (2C), 123.9 (2C), 16.4. *m/z* (ES+): 380.1 $[M+H^+]^+$

**(4-(4-Methylthiazol-5-yl)-2-nitrobenzyl)methanamine (11)**

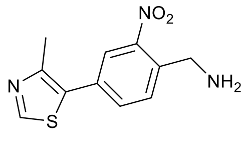

To a stirred suspension of **10** (1.70 g, 4.48 mmol) in ethanol (40 mL) under nitrogen was added hydrazine hydrate (0.65 mL, 13.45 mmol) in a dropwise fashion. The resulting reaction mixture was stirred at 80ºC for 3 hours. The reaction mixture was filtered and the filtrate was concentrated under reduced pressure to afford the title compound as an orange solid which was used directly in the next reaction without further purification (1.11 g, 99%). $^1$H NMR ($CDCl_3$): $\delta$ 8.75 (s, 1H), 8.07 (d, $J$ = 1.4 Hz, 1 H), 7.69-7.67 (d, 2 H), 4.16 (s, 2 H), 2.56 (s, 3 H). *m/z* (ES+): 250.1 $[M+H^+]^+$

***tert*-Butyl (4-(4-methylthiazol-5-yl)-2-nitrobenzyl)carbamate (12)**

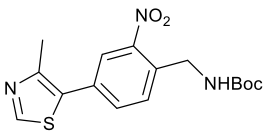

To a stirred solution of **11** (1.10 g, 4.42 mmol) and di-*tert*-butyl dicarbonate (1.20 g, 5.30 mmol) in THF (14 mL) under nitrogen was added triethylamine (1.85 mL, 13.26 mmol). The resulting reaction mixture was stirred for 1 hour at ambient temperature, treated with water (30 mL) and extracted with ethyl acetate (3 × 30 mL). The combined organic extracts were washed with brine (3 × 50 mL), dried over anhydrous magnesium sulfate and concentrated under reduced pressure to a red oil. Purification by flash column chromatography, eluting with 0-100% EtOAc/heptane, afforded the title compound as a yellow oil (1.10 g, 71%). $^{1}$H NMR ($CDCl_3$): $\delta$ 8.67 (s, 1H), 8.05 (bs, 1 H), 7.64-7.63 (m, 2 H), 5.57-5.56 (m, 1 H), 4.55 (d, $J$ = 6.3 Hz, 2 H), 2.48 (s, 3 H), 1.38 (s, 9H). $^{13}$C NMR ($CDCl_3$): $\delta$ = 155.9, 152.8, 151.4, 149.9, 148.2, 134.2, 132.7, 131.6, 130.9, 129.1, 125.3, 79.9, 42.1, 28.3, 16.1. *m/z* (ES+): 350.2 $[M+H^{+}]^{+}$

***tert*-Butyl (2-amino-4-(4-methylthiazol-5-yl)benzyl)carbamate (13)**

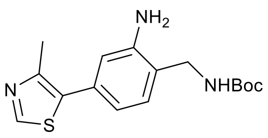

To a stirred solution of **12** (148 mg, 0.42 mmol) in MeOH (2 mL) was added zinc dust (277 mg, 4.24 mmoland ammonium formate (267 mg, 4.24 mmol), and the reaction mixture was stirred at ambient temperature for 30 min. After filtration through diatomaceous earth, the filtrate was concentrated under reduced pressure. Purification by flash column chromatography, eluting with 0-10% MeOH/DCM, afforded the title compound as an off-white solid (113 mg, 85%). $^{1}$H NMR (500 MHz, $CDCl_3$) $\delta$ 8.64 (s, 1H), 7.06 (s, 1H), 6.77 – 6.67 (m, 2H), 4.27– 4.24 (s, 2H), 2.52 (s, 3H), 1.45 (s, 9H). *m/z* (ES+): 320.0 $[M+H^{+}]^{+}$

***tert*-Butyl (*S*)-(1-(4-(4-methylthiazol-5-yl)phenyl)-2-oxoethyl)carbamate (14)**

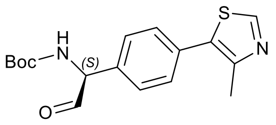

To a stirred solution of **1** (117 mg, 350 µmol) in DCM (2 mL) at 0 ºC was added Dess-Martin periodinane (297 mg, 700 µmol). The resulting reaction mixture was stirred at 0 ºC for 10 minutes, then at ambient temperature for 2 hours. The mixture was stirred vigorously with a saturated aqueous solution of sodium hydrogen carbonate containing 10% sodium thiosulfate (20 mL) for 10 minutes, and extracted with DCM (2 × 10 mL). The combined organic extracts were washed with brine (2 × 30 mL), dried over anhydrous magnesium sulfate and concentrated under reduced pressure to afford the title compound as a yellow solid, which was used directly in the next reaction without further purification (98 mg, 35%). $^{1}$H NMR ($CDCl_3$): $\delta$ 9.58 (s, 1 H), 8.71 (s, 1H), 7.50 (d, $J$ =

8.3 Hz, 2H), 7.39 (d, *J* = 8.3 Hz, 2H), 5.84 (bs, 1 H), 5.38 (bs, 1 H), 2.54 (s, 3 H), 1.45 (s, 9 H). *m/z* (ES+): 333.1 $[M+H^+]^+$

**_tert_-Butyl (_S_)-(2-((2-(((_tert_-butoxycarbonyl)amino)methyl)-5-(4-methylthiazol-5-yl)phenyl)amino)-1-(4-(4-methylthiazol-5-yl)phenyl)ethyl)carbamate (15)**

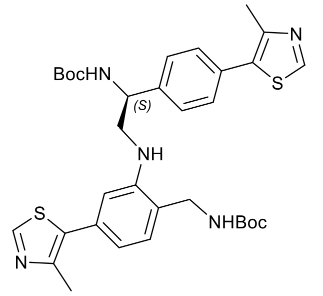

To a stirred solution of **14** (98 mg, 295 μmol) and **13** (94 mg, 295 μmol) in DCM (3 mL) was added sodium triacetoxyborohydride (93 mg, 443 μmol) in a portionwise fashion, and the resulting reaction mixture was stirred at ambient temperature under nitrogen for 24 hours. The reaction mixture was filtered through diatomaceous earth, treated with water and extracted with ethyl acetate. The combined organic extracts were washed with brine, dried over anhydrous magnesium sulfate and concentrated under reduced pressure to yield a yellow oil. Purification by flash column chromatography, eluting with 0-100% EtOAc/heptane, afforded the title compound as a yellow oil (24.7 mg, 17%). $^1$H NMR ($CDCl_3$) $\delta$ 8.69 (s, 1 H), 8.65 (s, 1 H), 7.43 (d, *J* = 2.5 Hz, 4H), 7.07 (d, *J* = 7.8 Hz, 1H), 6.85 (dd, *J* = 7.8 Hz, *J* = 1.5 Hz, 1H), 6.58 (d, *J* = 1.5 Hz, 1H), 5.44 (bs, 1 H), 5.06-5.00 (m, 1 H), 4.81 (d, *J* = 8.3 Hz, 1H), 4.40-4.21 (m, 1 H), 2.54 (s, 3 H), 2.52 (s, 3 H), 1.52 (s, 9 H). $^{13}$C NMR ($CDCl_3$) $\delta$ 171.2, 150.5, 150.1, 148.6, 148.3, 132.1, 131.8, 131.5, 131.5, 131.4, 130.1, 129.5, 127.5, 126.5, 126.3, 119.9, 118.9, 115.7, 60.4, 31.9, 28.4, 22.7, 21.1, 16.2, 16.2, 14.2. *m/z* (ES+): 535.1 $[M\text{-}Boc+H^+]^+$

**(2_S_,4_R_)-1-((_R_)-2-(1-Fluorocyclopropane-1-carboxamido)-3,3-dimethylbutanoyl)-N-(2-(((2_S_)-2-((1_R_,4_S_)-2-((_R_)-2-(1-fluorocyclopropane-1-carboxamido)-3,3-dimethylbutanoyl)-4-hydroxycyclopentane-1-carboxamido)-2-(4-(4-methylthiazol-5-yl)phenyl)ethyl)amino)-4-(4-methylthiazol-5-yl)benzyl)-4-hydroxypyrrolidine-2-carboxamide (ALS18)**

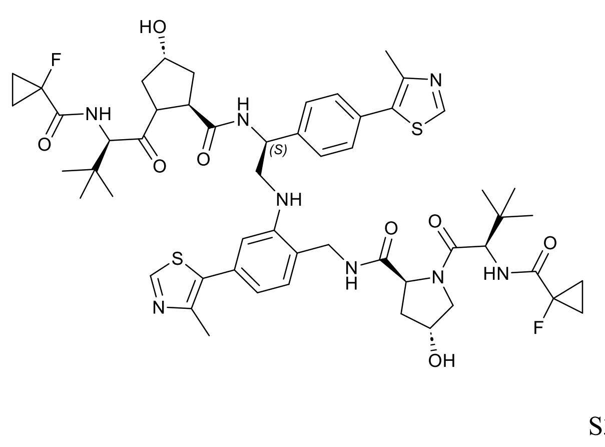

To a stirred solution of **15** (24.7 mg, 37 μmol) in DCM/MeOH (1:1) was added hydrogen chloride (1 mL, 4M solution in 1,4-dioxane). The resulting mixture was stirred at room temperature under nitrogen for 18 hours and then concentrated under reduced pressure to yield the Boc-deprotected intermediate as a yellow oil. The residue was redissolved in DMF (1 mL) and sequentially treated with (2*R*,4*R*)-1-((*R*)-2-(1-Fluorocyclopropane-1-carboxamido)-3,3-dimethylbutanoyl)-4-hydroxypyrrolidine-2-carboxylic acid, (16.6 mg, 50 μmol), DIPEA (17.5 μL, 100 μmol), HATU (21 mg, 50 μmol) and stirred at ambient temperature overnight. The reaction mixture was diluted with ethyl acetate and washed with brine (3 × 10 mL). The organic layer was dried over anhydrous sodium sulfate and concentrated under reduced pressure. Purification by preparative HPLC, eluting with 5-95% 0.1% formic acid in water/acetonitrile, afforded the title compound as a white solid (2.5 mg, 9%). $^{1}$H NMR (500 MHz, $CDCl_3$): δ 8.67 (s, 1H), 8.64 (s, 1H), 8.36 (d, *J* = 6.7 Hz, 1H), 7.91– 7.90 (m, 1H), 7.55 (d, *J* = 8.2 Hz, 2H), 7.39 (d, *J* = 8.2 Hz, 2H), 7.17 (d, *J* = 7.7 Hz, 1H), 6.99 – 6.90 (m, 2H), 6.71 (dd, *J* = 7.7, 1.7 Hz, 1H), 6.61 (s, 1H), 5.07 (d, *J* = 4.6 Hz, 1H), 4.89 (d, *J* = 5.0 Hz, 1H), 4.84 (t, *J* = 7.5 Hz, 1H), 4.63 (dd, *J* = 9.4, 7.1 Hz, 1H), 4.57 (d, *J* = 7.3 Hz, 1H), 4.56 – 4.45 (m, 4H), 4.43 (s, 1H), 4.35 (d, *J* = 7.8 Hz, 1H), 4.07 (dd, *J* = 14.6, 4.0 Hz, 1H), 3.94 (d, *J* = 11.3 Hz, 1H), 3.90 – 3.74 (m, 3H), 3.70 – 3.60 (m, 4H), 3.56 – 3.49 (m, 1H), 2.61 (s, 3H), 2.51 (s, 3H), 2.37 (ddd, *J* = 12.5, 7.7, 4.4 Hz, 1H), 2.22 (ddd, *J* = 13.4, 9.5, 4.1 Hz, 1H), 2.05 (dd, *J* = 13.3, 7.3 Hz, 1H), 1.96 (d, *J* = 11.8 Hz, 1H), 1.41 – 1.30 (m, 8H), 0.96 (s, 9H), 0.85 (s, 9H). HRMS (ES+) calculated for $C_{54}H_{68}F_2N_8O_8S_2$ 1060.3028 $[M+H^{+}]^{+}$, found 1060.4593

#### HPLC/MS Traces and $^1$H NMR for CC124, ALS14, ALS18

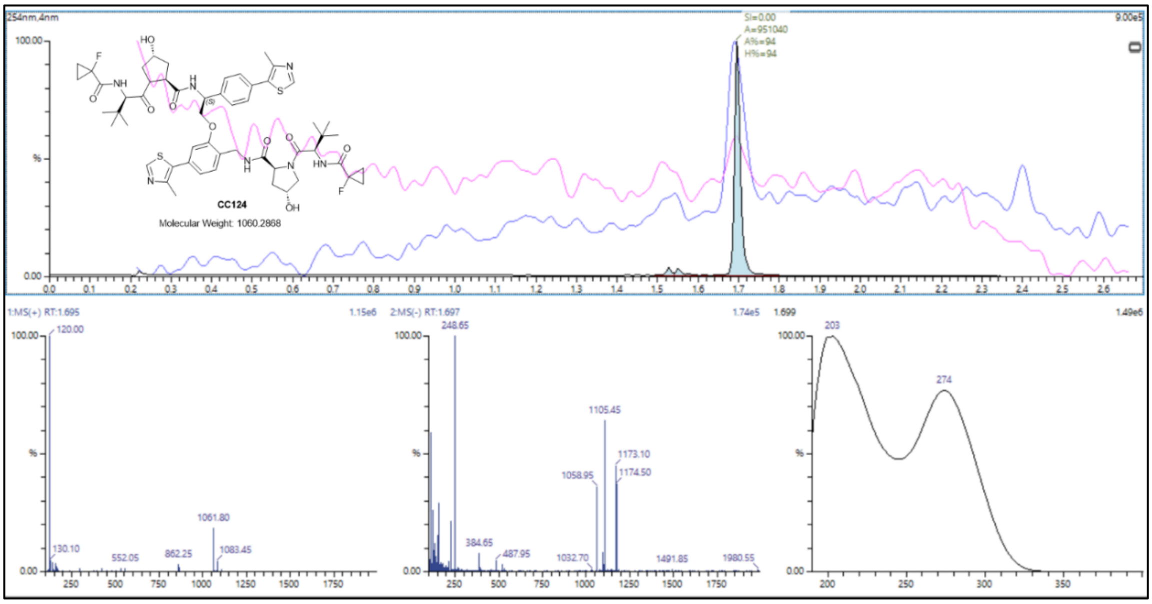

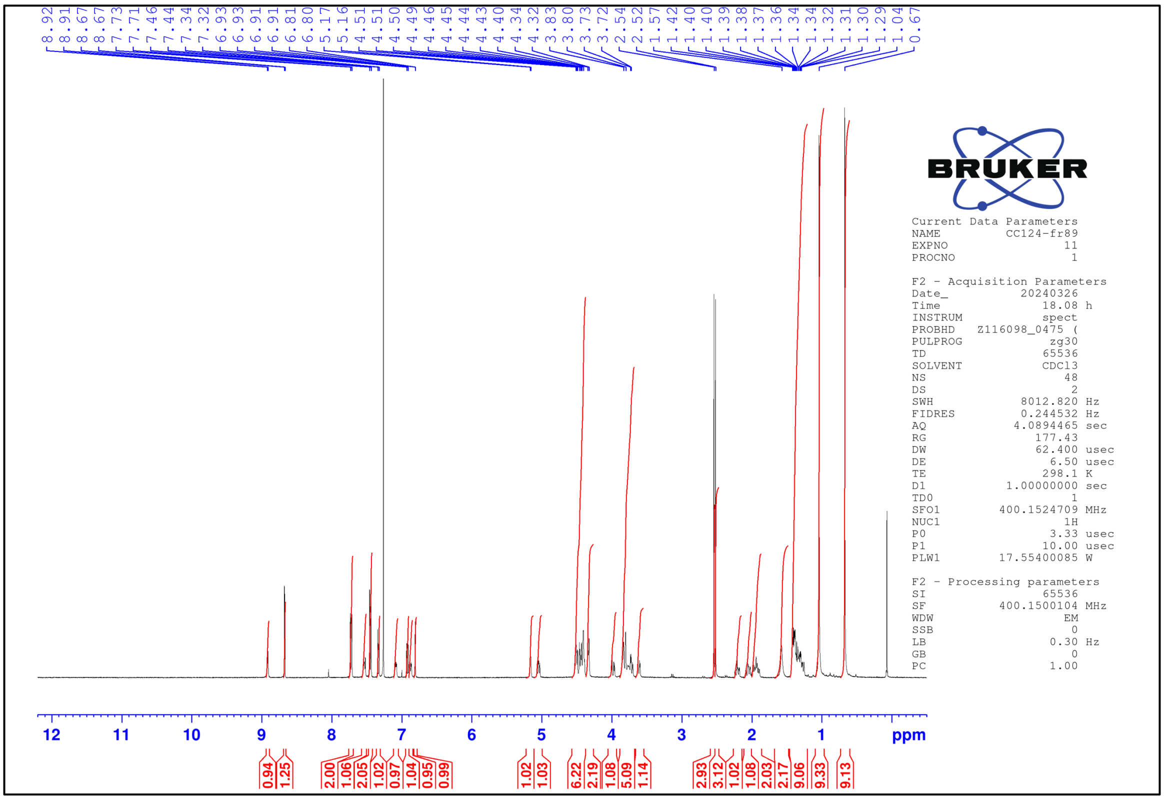

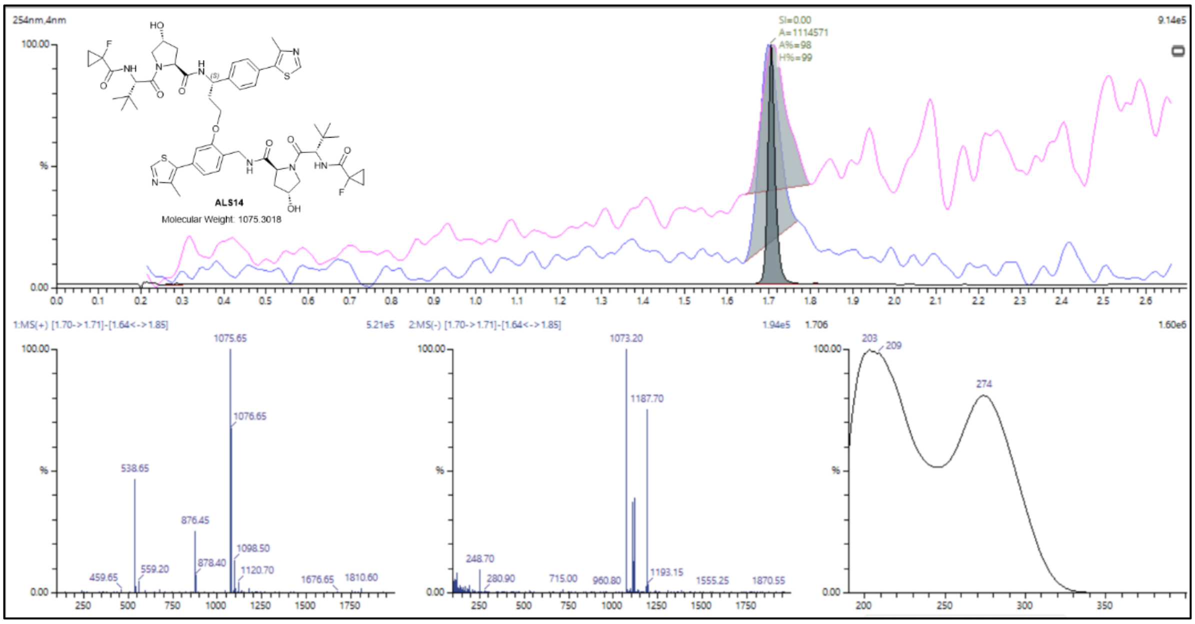
ALS14
Molecular Weight: 1075.3018

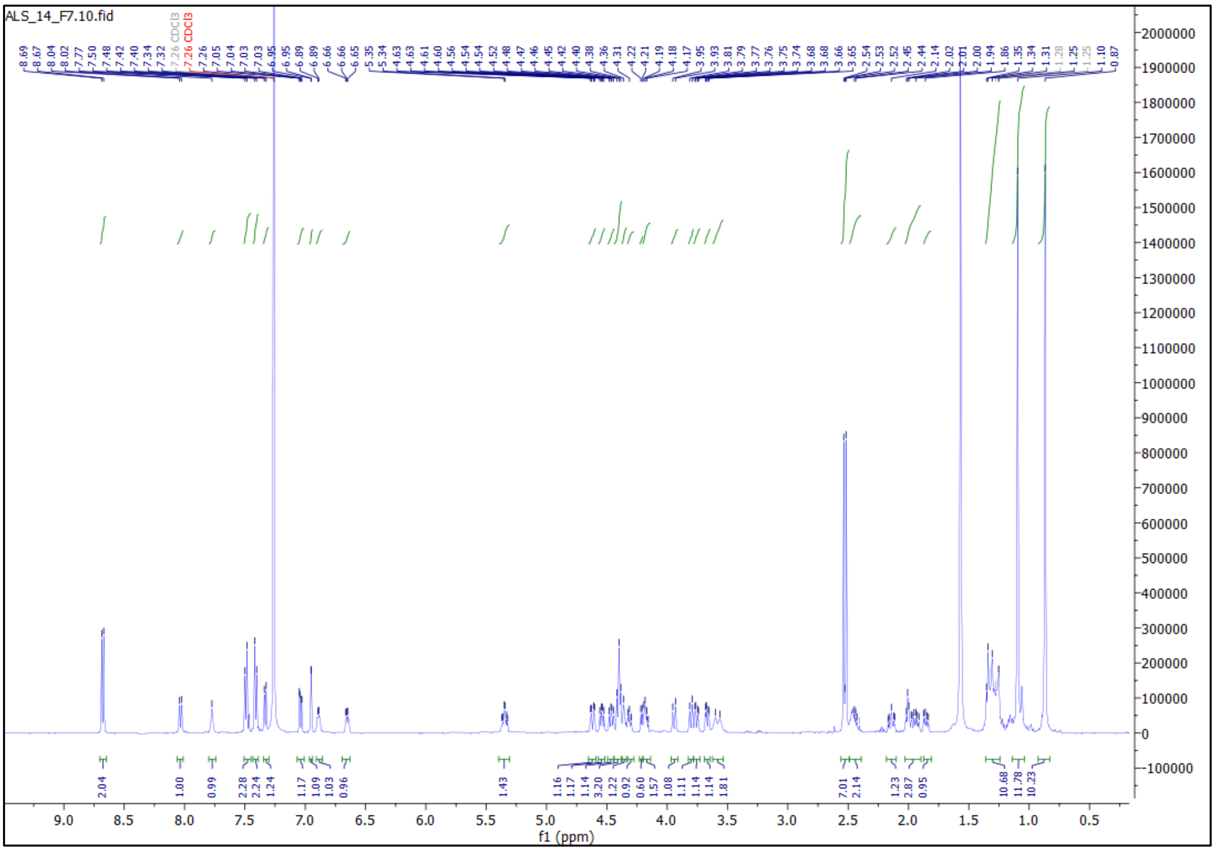
ALS_14_F7.10.fid
f1 (ppm)

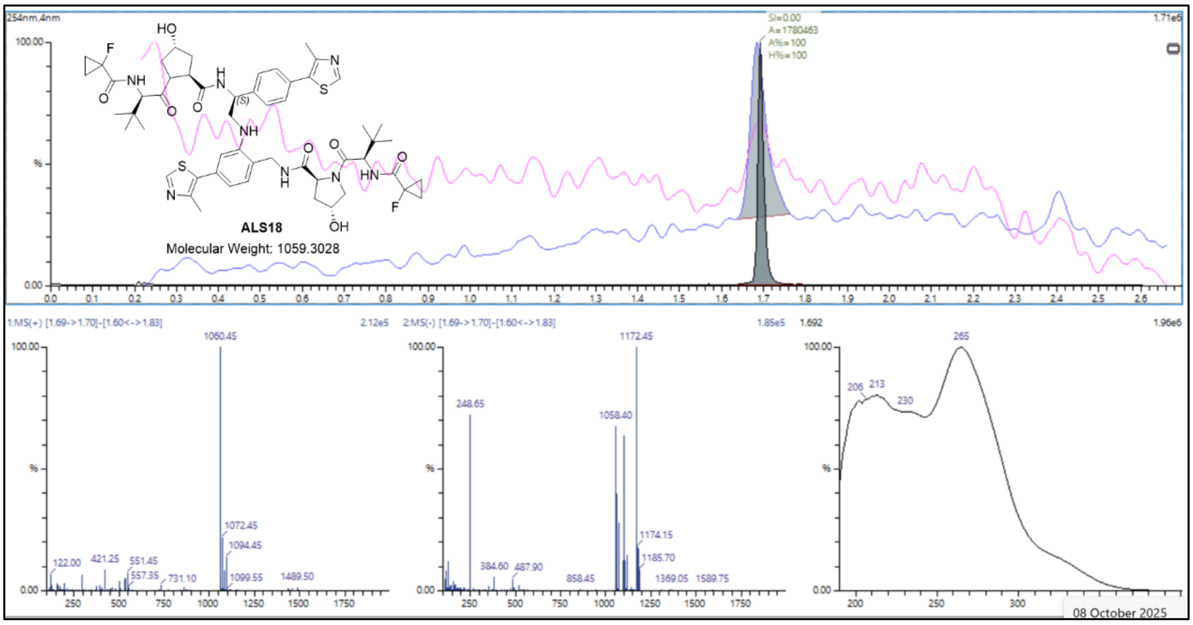
ALS18
Molecular Weight: 1059.3028
1060.45
1172.45
265
08 October 2025

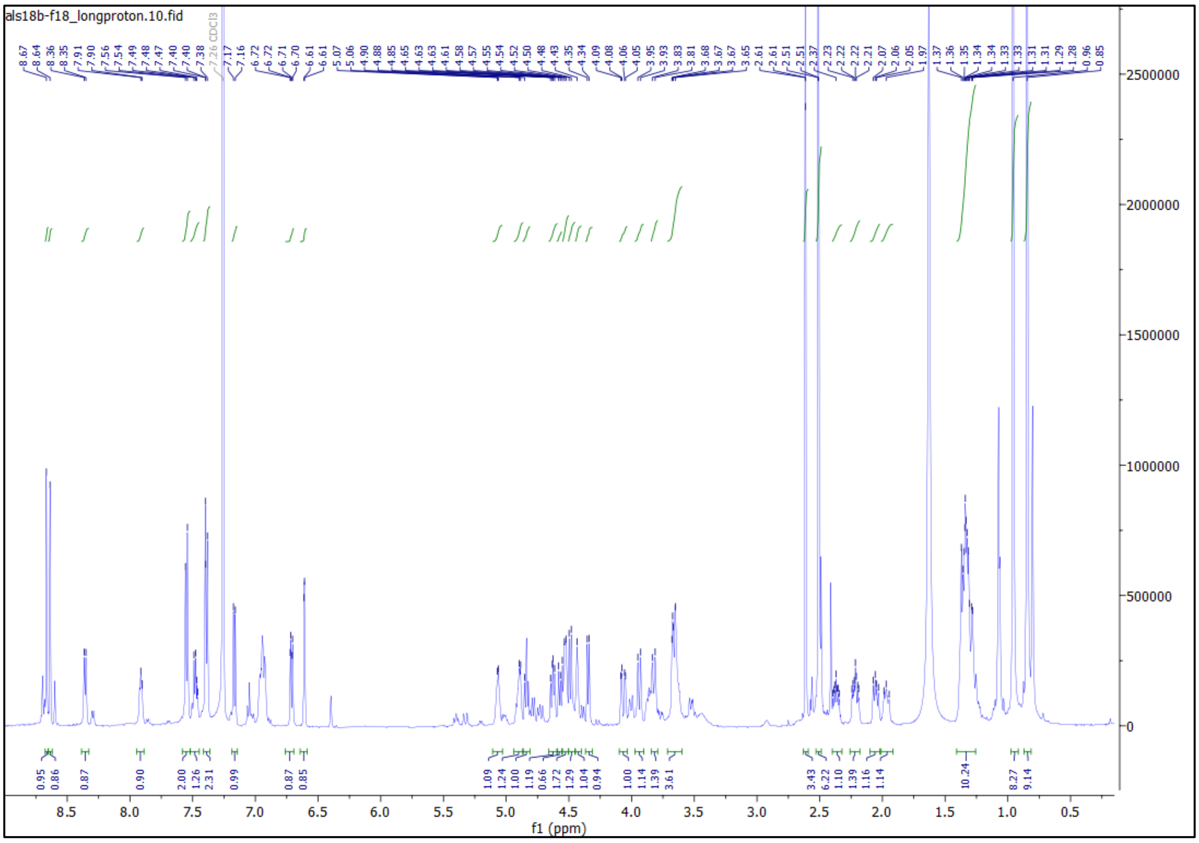
als18b-f18_longproton.10.fid
f1 (ppm)

### 2. Biology

#### 2.1 Protein expression and purification

Pure recombinant VCB (VHL residues 54-213, EloB residues 1-104 and EloC residues 17-112), NEDD8 (residues 1-76), APPBPA-UBA3, UBE2M, GACG-ubiquitin, UBE2D2, UBE2R1 and UBA1 were obtained from our previous study.[1]

For NEDD8-CRL2$^{VHL}$, single colonies of Cul2-Rbx1 cDNA[2] transformed into *E. coli* BL21(DE3) cells were grown overnight in LB supplemented with ampicillin (100 µg mL$^{-1}$) at 37 °C with shaking. The overnight culture was diluted 1:100 in LB supplemented with ampicillin (100 µg mL$^{-1}$) and grown to an optical density (OD600) of ~0.6. Protein expression was induced for 16 hours at 16 °C with 0.2 mM IPTG. The cells were harvested by centrifugation and frozen at -80 °C as pellets until further purification. The cells were thawed, resuspended in 30 mM Tris-HCl, 200 mM NaCl, 5 mM DTT, pH 7.5, supplemented with 5 mM $MgCl_2$, 1 µg/mL DNAse I and 1X EDTA-free Roche protease inhibitor cocktail, and lysed by cell disruption at 30 kpsi. Cellular debris was removed by centrifugation. Cul2-Rbx1 was purified on a 5 mL HisTrap™ FF Ni NTA affinity column (Cytiva) and eluted with an imidazole concentration gradient from 0 to 300 mM. The protein was desalted into 20 mM HEPES, 150 mM NaCl, 0.5 mM TCEP, 5% (v/v) glycerol, pH 7.5 with a HiPrep 26/10 desalting column (Cytiva) and incubated with TEV protease and an excess of pure recombinant VHL-EloB-EloC. VHL-EloB-EloC-Cul2-Rbx1 (CRL2$^{VHL}$) was bound to a 5 mL StrepTrap XT column (Cytiva), washing with 100 mM Tris-HCl, 150 mM NaCl, pH 8.0 and eluting in 100 mM Tris-HCl, 300 mM NaCl, 50 mM biotin and 5% (v/v) glycerol, pH 8.0. Purified CRL2$^{VHL}$ (1.7 µM) was incubated with pure recombinant APPBP1-UBA3 (250 nM), UBE2M (1.2 µM), NEDD8 (20 µM), ATP (1 mM) and $MgCl_2$ (5 mM) for 10 minutes at 37 °C to allow the neddylation reaction to occur. NEDD8-CRL2$^{VHL}$ was finally purified by size exclusion chromatography on a HiLoad™ 16/600 Superdex™ 200 pg column (Cytiva) in 20 mM HEPES, 150 mM NaCl, 0.5 mM TCEP, 5% (v/v) glycerol, pH 7.5, concentrated to 3.2 or 6 µM, then flash frozen with liquid nitrogen and stored at -80 °C.

#### 2.2 Ubiquitin labelling with maleimide Alexa Fluor 488

Alexa Fluor 488 C5 maleimide dye was dissolved to 5 mM in anhydrous DMSO. Ubiquitin cysteine mutant recombinant protein was buffer exchange into 20 mM HEPES, 150 mM NaCl, pH 7.0 using a 10/300 GL Superdex 75 Increase prepacked column (Cytiva). The fractions containing the protein were combined, concentrated to 100 μM and labelled with a 5 times molar excess of fluorescent dye for 2 hours at room temperature. The excess dye was removed by gel filtration on a 10/300 GL Superdex 75 Increase prepacked column (Cytiva), eluting in 20 mM HEPES, 150 mM NaCl, 0.5 mM TCEP, pH 7.5. The dye and subsequently dye-labelled protein were protected from light throughout, flash frozen and stored at -80 °C.

#### 2.3 In vitro ubiquitination assay and immunoblotting

The E1 Ube1 (150 nM), the UBE2D2 or UBE2R1 (5 μM), $CRL2^{VHL}$ or NEDD8-$CRL2^{VHL}$ (2 μM), and homoPROTAC (5 μM) were mixed with ubiquitin (98 μM) and Alexa Fluo 488-labelled ubiquitin (2 μM) and incubated in 20 mM HEPES, 150 mM NaCl, 0.5 mM TCEP, 5 mM $MgCl_2$, pH = 7.5 for 5 minutes at room temperature. ATP (3 mM) was added, and the mixture was incubated at room temperature overnight. Samples were quenched by diluting 1:3 in 2X reducing LDS sample buffer (NuPAGE) and proteins were separated by SDS-PAGE on a 12% Bis-Tris gel (NuPAGE) in MOPS running buffer, run at 185 V for 50 min. Proteins were transferred to nitrocellulose membrane using iBlot 2 Gel Transfer Device (Invitrogen). Membranes were blocked for 1 hour in 5% milk in TBS-T and incubated overnight with primary antibodies, then for 1 hour with secondary antibodies and dyes (1:5000) before imaging on the ChemiDoc (Bio-Rad).

#### 2.4 Electrophoresis Mobility Shift Assay

10% polyacrylamide gels were prepared from 40% acrylamide solution (3.75 mL), $ddH_2O$ (7.5 mL), 1.5 M Tris buffer (3.75 mL, pH = 8.8), 10% (*w/v*) aqueous ammonium persulfate solution (150 μL) and tetramethyl ethylenediamine (10 μL) and were polymerised in washed, SDS-free gel casting system for 1 h and used within one week. VCB recombinant protein was diluted in sample buffer (20 mM HEPES, 150 mM NaCl, pH adjusted to pH = 7.0) to a final sample concentration of 12.11 μM and 0.5 μL of 20X EtOH stocks of the respective PROTACs were added (final concentration 24.22 μM) and were incubated for 30 min at room temperature. 1 μL of native gel dye (made from 2.5 mL tris-glycine running buffer, 5 mL glycerol, 1 mL bromophenol blue (1%) and 1.5 mL ddH2O) were added, and the samples were loaded into the native gel and resolved at 100 V for at least 120 min on ice in Tris-glycine running buffer (25 mM Tris base and 192 mM glycine). Protein bands were visualised with Instant Blue Coomassie stain.

##### 2.5 Preparation of samples for cryo-electron microscopy

Frozen stocks of recombinant protein and homoPROTAC were thawed and kept at 4 °C throughout the sample preparation process. 6 μM NEDD8-CRL2$^{VHL}$ (1 equivalent) and CM11 (10 equivalents) were incubated for 10 minutes at 4 °C. The complexes were desalted on a 0.5 mL 7K MWCO Zeba™ Spin Desalting Column in 20 mM HEPES, 150 mM NaCl, 0.5 mM TCEP, pH 7.5. Quantifoil R1.2/1.3 holey carbon copper 400 mesh grids were glow discharged for 60 seconds at 35 mA using a Quorum SC7620. 3.5 μL of protein at approximately 4 μM was applied to the cryo-EM grids and was vitrified in liquid ethane on a Vitrobot Mark IV (Thermo Fisher Scientific) at 4 °C and 100% humidity (wait time = 10 s, blot force = 4, blot time = 3.5 s, blot total = 1, drain time = 0 s).

##### 2.6 Cryo-electron microscopy data acquisition

Cryo-EM data were collected on Glacios transmission electron microscope (Thermo Fisher Scientific) operating at 200 keV. Micrographs were acquired using a Falcon4i direct electron detector (Thermo Fisher Scientific), operated in electron counting mode. A total electron exposure of 57 $e^-/Å^2$ was applied. EPU (Thermo Fisher Scientific, version 3.0) was used to collect micrographs at 190,000k nominal magnification (0.74 Å/pixel at the specimen level) with a nominal defocus range of -1.7 to -3.2 μm. Stage shifts with aberration-free image shift (AFIS) mode was used to centre multiple foil holes and image shift was used to acquire high magnification images in the centre of each targeted hole. 7,216 movies were collected in EER format.

##### 2.7 Cryo-electron microscopy image analysis and model building

Image processing pipelines are described in Appendix Figures S1 and S2. Cryo-EM movies were imported into CryoSPARC v.4.4.0-v4.4.1[3] for patch motion correction, patch CTF estimation and manual curation. Particle picking was either performed manually, or performed with crYOLO using a general model for low-pass filtered images.[4] Particles were submitted to 2D classification and good templates were used for template picking. Particles were extract ed with a 600-pixel box size (3x binning). Classification was achieved 2D classification, and 3D reconstruction was performed with *ab initio* reconstruction and non-uniform refinement in CryoSPARC.[5] 15 cryo-EM maps from independent refinement jobs with both C1 and C2 symmetry were obtained following the above methodology. Representative 2D classes, orientation diagnostics, local resolution estimations and gold-standard Fourier shell correlation (GSFSC) curves were generated with CryoSPARC[5]. Model building was achieved using atomic models from AlphaFold and PDB entries 5N4W, 8RX0, 5NVX, which were docked into the cryo-EM maps using rigid-body fitting with UCSF ChimeraX.[6] The models were refined using ISOLDE[7] and Phenix until reasonable agreement between the model and data were achieved.

##### 2.8 Mass photometry

Interferometric scattering microscopy was carried out using the oneMP (Refeyn). Gaskets wells (Grace Bio-labs CW-50R-1.0) along with high precision 24 x 50 mm coverslips (Marienfeld) were used. 10 µL of buffer (20 mM HEPES, 150 mM NaCl, 0.5 mM TCEP, pH 7.5, filtered through a 0.22 µm syringe filter) was used for focus in regular mode (128 x 34 binned pixels 18.0 $\mu m^2$ detection area). After focus was locked with buffer, 10 µL of sample containing 50 nM (NEDD8)-CRL2$^{VHL}$ (1 equivalent) and DMSO or homoPROTAC (2 equivalents) was added and mixed by pipetting. The data was recorded for 60 seconds using AquireMP software (Refeyn). Calibration was performed using the MassFerence® P1 calibration standard (Refeyn). The data was processed and analyzed in DiscoverMP (Refeyn).

##### 2.9 Protein conjugation to switchSENSE® ssDNA

ssDNA *cNL-A48* (from coupling kit *HK-NHS-1*, Dynamic Biosensors) and *cNL-B48* (from coupling kit *HK-NHS-4*, Dynamic Biosensors) were dissolved in 50 mM $Na_2HPO_4$/$NaH_2PO_4$, 150 mM NaCl, pH 7.2, incubated with crosslinker (Dynamic Biosensors) at room temperature for 20 minutes, then desalted into 50 mM $Na_2HPO_4/NaH_2PO_4$, 150 mM NaCl, pH 8.0. 200 µg of VCB recombinant protein was incubated with the ssDNA at room temperature for 1 hour. Excess DNA and unreacted protein were removed by ion exchange chromatography on a *PF-CC-1-1* chromatographic column (Dynamic Biosensors) using proFIRE® FPLC system (Dynamic Biosensors). The fractions containing pure protein-ssDNA conjugates were pooled, concentrated and buffer exchanged into 10 mM $Na_2HPO_4/NaH_2PO_4$, 40 mM NaCl, 0.05 % Tween 20, 50 µM EDTA, 50 µM EGTA, pH 7.4, before being flash frozen and stored at -80 °C. Protein-ssDNA concentration was estimated using a Nanodrop Microliter UV/Vis spectrophotometer (Thermo Fisher Scientific). The absorbance at 260 nm was measured and the concentration of protein-ssDNA was calculated following the Beer-Lambert equation using the molar extinction coefficient of 490,000 L $mol^{-1}$ $cm^{-1}$ as given by Dynamic Biosensors.

#### 2.10 heliX® switchSENSE® binary binding assay

SwitchSENSE® Adapter Strand measurements were performed on a dual-colour $heliX^+$ instrument using a heliX® Adapter Biochip (*ADP-48-2-0*, Dynamic Biosensors). 100 nM of Adapter Strand 1 with red dye Ra (for VCB) or 100 nM of Adapter Strand 1 with green dye Ga (for BET bromodomains) was incubated with 120 nM of VCB-ssDNA at room temperature for 20 minutes with shaking (500 rpm). 100 nM of Adapter Strand 2 with red dye Ra prehybridised with ligand-free strand was added and the mixture was used to functionalise the biochip surface directly before PROTAC measurements.

Interaction analysis was performed in fluorescence proximity sensing (FPS) mode. 10 mM stocks of PROTACs in DMSO were diluted in 10 mM HEPES, 140 mM NaCl, 0.05 % Tween20, 50 µM EDTA, 50 µM EGTA, pH 7.4 to final concentrations of 125 nM, 25 nM, 5 nM, 1 nM and 200 pM. Measurements for the association with PROTACs (2 minutes) and dissociation with buffer (10 mM HEPES, 140 mM NaCl, 0.05 % Tween20, 50 µM EDTA, 50 µM EGTA, pH 7.4) were performed at a constant flow rate of 200 µL/min. The Y-structure was excited using an LED power setting of 2 and the fluorescence signal was monitored. The biochip was functionalised only once for each PROTAC as full dissociation from its binary binding partner was observed after 15 minutes of dissociation. Biochip regeneration was performed using the regeneration buffer *HK-REG-1* (Dynamic Biosensors). The PROTAC samples were maintained at 15 °C and the biochips were maintained at 25 °C throughout the experiments. Experiment design, workflow and data analysis were performed with heliOS

2023.1 software (Dynamic Biosensors). The association and dissociation rates ($k_{on}$ and $k_{off}$) and dissociation constants ($K_D$) were derived from a global single exponential fit model after blank referencing correction and reference electrode correction.

#### 2.11 heliX® switchSENSE® ternary complex formation assay

SwitchSENSE® Y-structure measurements were performed on a dual-colour heliX$^+$ instrument using a heliX® Adapter Biochip (*ADP-48-2-0*, Dynamic Biosensors). 100 nM of Y-structure Adapter with red dye Ra, 100 nM of Y-structure Adapter with green dye Ga (Dynamic Biosensors), 120 nM of VCB-ssDNA was incubated at room temperature for 2 hours with shaking (400 rpm). 100 nM of cAnchor 2 (Dynamic Biosensors) was added and the mixture was used to functionalise the biochip surface directly before PROTAC measurements. Interaction analysis was performed in fluorescence proximity sensing (FPS) mode. 10 mM stocks of PROTACs in DMSO were diluted in 10 mM HEPES, 140 mM NaCl, 0.05 % Tween20, 50 µM EDTA, 50 µM EGTA, pH 7.4. Measurements for the association with PROTACs (2 minutes) and dissociation with buffer (10 mM HEPES, 140 mM NaCl, 0.05 % Tween20, 50 µM EDTA, 50 µM EGTA, pH 7.4) were performed at a constant flow rate of 500 µL/min. The Y-structure was excited using an LED power setting of 2 and the FRET signal was monitored. The biochip was regenerated and freshly functionalised for each. Biochip regeneration was performed using the regeneration buffer *HK-REG-1* (Dynamic Biosensors). The PROTAC samples were maintained at 15 °C and the biochips were maintained at 25 °C throughout the experiments. Experiment design, workflow and data analysis were performed with heliOS 2023.1 software (Dynamic Biosensors). The association and dissociation rates ($k_{on}$ and $k_{off}$) and dissociation constants ($K_D$) were derived from a global single exponential fit model after blank referencing correction.

#### 2.12 Mammalian cell culture

HEK293 cell lines, originally sourced from ATCC, were provided by the MRC PPU reagents facility at the University of Dundee. HEK293 cells were cultured in DMEM (Gibco) supplemented with 10% fetal bovine serum (FBS; Thermo Fisher), 100 U ml$^{-1}$ penicillin-streptomycin (Thermo Fisher) and 2 mM L-glutamine (Thermo Fisher). All cell lines were grown in a humidified incubator at 37 °C and 5 % $CO_2$ and routinely tested for mycoplasma contamination.

#### 2.13 Cellular target engagement assay

HEK 293 cells (20 mL) were transfected at a density of 2 × $10^5$ cells/mL with 1 µg VHL-NanoLuc® fusion DNA (Promega), 9 µg Transfection Carrier DNA (Promega) and 30 µL FuGENE® HD in 1 mL of Opti-MEM growth medium and incubated for 20 hours at 37 °C and 5% $CO_2$. The transfected cells were pelleted, washed, and resuspended in Opti-MEM to a density of 2.2 × $10^5$ cells/mL. The cells were seeded in a white 96-well plate (Corning), treated with PROTAC in DMSO solution, a solution NanoBRET™ Tracer (Promega) to a final concentration of 125 nM and 500 µL/mL digitonin, and incubated for 10 minutes. Directly prior to measurement, the cells were treated with 50 µL of a solution of NanoBRET™ Detection Reagent (1:400 dilution) in Opti-MEM. The plates were mixed on an orbital shaker for 30 seconds at 500 rpm at room temperature and read on a PHERAstar FSC microplate reader (BMG Labtech) with measurements at a donor emission wavelength of 460 nm and an acceptor emission wavelength of 618 nm. BRET values in milliBRET units (mBU) were obtained for each PROTAC by dividing the acceptor emission value (610 nm) by the donor emission value (450 nm) for each sample. BRET values versus PROTAC concentration were plotted on GraphPad Prism 9 and the $EC_{50}$ was obtained by fitting a nonlinear regression curve of 4-parameter variable slope for each PROTAC.

#### 2.14 Cell treatment for immunoblotting

HEK 293 cells were plated in 6-well plates at varying densities (4-5 × 105 cells/mL) 24 h before treatment depending on the experimental setup. Cells were treated in fresh medium with the indicated compounds under indicated conditions with a final DMSO concentration of 0.5% (v/v). After compound treatment, the medium was removed, and the cells were washed with ice-cold Phosphate-Buffered Saline (PBS) and lysed on ice with 100 µL RIPA lysis and extraction buffer (Thermo Fisher Scientific, #89900) supplemented with complete EDTA-free protease inhibitor cocktail (11873580001, Roche). Cells were incubated for 15 min on ice and then detached from the surface by scraping. After removal of the insoluble fraction by centrifugation at 15,000 g at 4 °C for 15 min, supernatants were stored at −80°C. Protein concentration was determined by bicinchoninic acid (BCA) assay (Thermo Fisher Scientific, #23225) Cell lysates containing a quarter of a volume of 4× NuPAGE LDS sample buffer

(NP0007) supplemented with 10% DTT were heated at 95 °C for 5 min. Samples (20 to 30 µg) were loaded onto precast 4 – 12% bis–tris midi 20W or 26W gels (Thermo Fisher Scientific) and resolved at 90 V for 10 min and then at 130 V for 1.5 h with a NuPAGE MOPS SDS running buffer (Thermo Fisher Scientific). Gels are transferred onto nitrocellulose membranes using an iBlot3 system (Thermo Fisher Scientific). The transferred membrane was blocked with 5% (w/v) skim milk powder dissolved in tris-buffered saline with Tween (TBS-T) (50 mM tris base, 150 mM sodium chloride (NaCl), 0.1% (v/v) Tween-20) at room temperature for 1 h. Western blot images were obtained through detection with anti-VHL (Cell Signaling Technology, #68547; 1:1000), anti-HIF-1α (BD Biosciences, #610959, clone 54, 1:1000), anti-OH-HIF-1α (Pro564) (Cell Signaling Technology; #3434, 1:1000) and anti-Cul2 (Abcam, ab166917, 1:1000). Following overnight incubation with the primary antibodies at 4 °C, the membranes were washed two times for 10 min with TBS-T and then incubated with secondary antibodies anti-rabbit IRDye 800CW (LiCOR, 926-32213, 1:5000), anti-rabbit IRDye 680RD (LiCOR, 926-68073, 1:5000) anti-mouse 800CW (LiCOR, 926-32212, 1:5000) and hFAB™ rhodamine anti-tubulin antibody (BioRad, 12004165, 1:10,000) or 1 h at room temperature and protected from light. Thereafter, the membranes were washed with TBS-T three times for 10 min, and protein bands were acquired using a ChemiDoc MP imaging system (Bio-Rad). Band quantification was performed using Image Lab software and reported as ratio of each protein band relative to the lane's loading control. The values obtained were then normalised to vehicle control.
